# ONE-STEP tagging: a versatile method for rapid site-specific integration by simultaneous reagent delivery

**DOI:** 10.1101/2024.08.29.610246

**Authors:** Valentina Migliori, Michaela B. Bruntraeger, Ivan S. Gyulev, Florence Lichou, Thomas Burgold, Daniel P Gitterman, Sho Iwama, Andrew L. Trinh, Sam J. Washer, Carla P Jones, Gosia Trynka, Andrew R. Bassett

**Affiliations:** Wellcome Sanger Institute, Wellcome Genome Campus, Hinxton, Cambridge, CB10 1SA, UK

## Abstract

We present a novel, versatile genome editing method termed ONE-STEP tagging, which combines CRISPR-Cas9-mediated targeting with Bxb1 integrase-based site-specific integration for efficient, precise, and scalable protein tagging. Applied in hiPSCs, cancer cells, and primary T cells, this system enables rapid generation of endogenously tagged cell lines. By enhancing the nuclear localization signal (NLS) of the catalytically superior eeBxb1 integrase and co-delivering a DNA-PK inhibitor (AZD-7648), we achieved up to ∼90% integration efficiency at the ACTR10 locus. ONE-STEP tagging is robust across diverse loci and cell types and supports large DNA cargo integration, with efficiencies reaching 16.6% for a 14.4 kb construct. The method also enables multiplexed tagging of multiple proteins within the same cell and allows simultaneous CRISPR-based editing at secondary loci, such as gene knockouts or homology-directed repair (HDR) insertions. Importantly, we demonstrate successful application in primary T cells by targeting the T cell receptor (TCR) locus while simultaneously knocking out B2M, a key step toward generating immune-evasive, off-the-shelf CAR-T cells. Additionally, we introduce a dual-cassette version of the method compatible with universal donor plasmids, allowing use of entirely off-the-shelf reagents. Together, these advances establish ONE-STEP tagging as a powerful tool for both basic and therapeutic genome engineering.

## Introduction

Recent advancements in genome-editing technologies have provided efficient tools for specific genome modifications across various cell types and organisms. An important component of genome engineering is site-specific integration of DNA sequences. This allows genomic tagging of proteins to investigate their localisation, temporal dynamics and protein interactions. It also permits expression of transgenes under endogenous regulatory elements or at specific safe harbour loci for controlling transgene expression levels that is also important in cell therapeutics such as chimeric antigen receptor T cells (CAR-T) (1).

CRISPR-Cas9 enhanced homology-directed repair (HDR) has become a key technology for transgene integration (2). This involves creating a double-strand break (DSB), and supplying an excess of a homologous template DNA which is used to elicit a highly precise repair (3).

Short synthetic single-stranded DNA oligonucleotides (ssODN) with ∼100 nt of homology have been highly effective in many cell types (4, 5) but they are limited in cargo capacity to around 100-200 nt making insertion of large transgenes impossible. Longer plasmid DNA with around 500-1000 nt of homology can be used to integrate larger fragments (6), but there have been observations of complex, multimeric integration events at the on target site (7) as well as off-target integration of the plasmid DNA.

Others have used long ssODN (8) or linear double-strand DNA oligonucletides (dsODN) to circumvent these problems (9), but the former is difficult to produce, especially with longer cargos, and the latter is prone to random integration at off-target sites in the genome (10, 11). Protection of linear dsDNA with DNA structures or chemical modifications such as biotin has been shown to reduce non-specific integration and achieve tagging in some cell types (12, 13).

However, with all of these methods, it is still necessary to produce a template DNA for HDR that is different for every targeted site. Also, the efficiency of integration drops rapidly with increasing insertion size, making it difficult to insert inserts of more than 5-10 kb (14).

To overcome the size limitation, some groups have successfully combined HDR-mediated integration of a landing pad at a specific genomic locus followed by a second step of site-specific recombination with Cre (15), Bxb1 (16, 17) and Bxb1 compared to 14 other recombinases (18). The serine integrase Bxb1 is particularly useful for this, as it efficiently and specifically recombines heterologous attB and attP sites without known pseudo-sites in the human genome (19). Unlike Cre, its recombination is directional and irreversible (20), in this respect is is similar to phiC31 which has been used to insert large fragments in a number of vertebrate genomes (21), however phiC31 has a number of pseudo-sites in the human genome whereas Bxb1 does not (22). Recently, novel integrases with purported higher efficiencies in human cells than wild-type Bxb1 have been discovered such as Pa01 (23) but they have not been so extensively characterised.

However, such integration typically involves a two-step process, with clonal selection after the HDR event, making it quite lengthy and difficult to scale to multiple sites.

An alternative method for integration employs the non-homologous end joining (NHEJ) repair mechanism that ligates two dsDNA ends together (24, 25). By simultaneously cutting the genome and a donor DNA within the cell, this can be exploited to insert the donor DNA into any desired genomic site. This allows common donor plasmids to be used, avoiding the need for cloning, and making scaling of this method possible. It also has less of a length limitation than HDR-based methods, and tens of kilobases can be integrated using these methods.

However, there is no control over orientation of the insertion, in some cases the whole plasmid will be integrated, and the NHEJ repair mechanism can sometimes introduce small insertions and deletions around the genomic cut site.

Prime editing (PE) is another alternative for site specific integration that uses a modified single-guide RNA (pegRNA) containing the template for the desired edit, with a reverse transcriptase (RT) fused to Cas9 nickase (26). This makes a nick at a genomic locus and extends the genomic DNA by reverse transcription of the pegRNA to introduce the edit, and through manipulation of mismatch repair (MMR) pathways can be biased towards incorporation of the newly edited strand (27). PE offers advantages over HDR such as fewer mutagenic DSBs, but it also has limitations, including fewer targetable sites and limited insertion length (<50 bp) (28).

Recently, insertion of a site-specific recombinase site using PE, and simultaneous delivery of the cognate recombinase resulted in efficient integration of large cargos into specific genomic sites in a single-delivery reaction (PASTE technology) (29). Recombination efficiency has recently been improved using evolved Bxb1 integrase (eeBxb1 and evoBxb1) (30).

Similarly, template-jumping prime editing (TJ PE) has allowed insertions up to 800 bp (31). However, despite progress with computational prediction tools, it is still difficult to design effective PE guides without testing several permutations (32). Also, the target sites are somewhat limited by directionality of the PE process and availability of Cas9 cut sites (33).

All of these methodologies rely on the endogenous DNA repair pathways of HDR, NHEJ or MMR. The efficiency of these repair pathways varies significantly between cell types. Embryonic stem cells and induced pluripotent stem cells tend to favour HDR pathways (34), but many cancer cell lines and terminally differentiated cells preferentially repair through NHEJ (35, 36). Thus, the choice of the method will depend on the cell type and respective repair pathways that are active.

We present ONE-STEP tagging, a technology allowing simple, efficient, and directional integration of large transgenes at any genomic site using a single-step delivery protocol that employs single-stranded oligodeoxynucleotides (ssODN)-templated HDR combined with Bxb1-mediated integration. We show its utility in tagging at multiple genomic locations in multiple cell types including pluripotent stem cells, cancer cell lines and primary T cells. We further demonstrate that large cargos of up to 14.4 kb can be integrated. Tagging plus other Cas9-mediated knockout or HDR events can be performed simultaneously at two different sites. We further demonstrate its translational potential in primary human T cells, where we precisely integrated a 4.4 kb construct at the *TRAC* locus while simultaneously knocking out the immune-evasive gene B2M. This suggests that our method could be used to generate universal, off-the-shelf CAR-T cells. We also optimised the use of heterotypic Bxb1 recombinase sites to avoid the integration of plasmid backbones and enable dual-cassette tagging. Our system allows the use of completely off-the-shelf reagents, namely commercially available Cas9 protein, synthetic sgRNAs and ssODNs to define the genomic location and a common set of cargo vectors to define the inserted fragment which will make this methodology scalable to a large number of target sites.

## Materials and Methods

### ssODN design

ssODNs that contain a single recombination site use an attP variant sequence that is 58 bp long. ssODNs that contain two recombination sites use core attP variant sequences that are 48 bp long (derived from the 58bp variant). When convenient, the length of one of the two 48 bp long sites in the dual recombination cassette is extended by including an additional 5 bp from the 58 bp variant as described below. In this study, the recombination sites used in each part of the doublet pair of attP-attB sites contain different central dinucleotide mutants (heterotypic sites) in order to ensure the efficient integration of only the insert portion and not the backbone of the donor vector. This theoretically improves recombination efficiency by 50%. Supplementary Table S1 contains a breakdown of the recombination sites used throughout this work - attP, attB, resulting attR and attL sites and the lengths of DNA spacers used to keep insertions in-frame in the case of gene tagging. Due to the order of insertion of recombination sites in the genome - attP(s) being inserted in the genome first, and subsequently being recombined with a donor molecule containing attB(s) - the resulting recombinant sites and cassette are of the following structure: attR-insert-attL (this is the case for both the single and double pair of attP-attB site systems). In the case of 5’ end tagging (where an N-terminal protein fusion is desired), either the single or dual attP cassettes are designed to replace the start codon of the targeted CDS. In the case of 3’ end tagging (where a C-terminal protein fusion is desired), either the single or dual attP cassettes are designed to replace the stop codon of the targeted CDS.The single attP-attB pair results in attR and attL sites that are 52 bp long post-integration. The double attP-attB pair results in attR and attL sites which are 47 bp long post-integration. This difference necessitates a slightly different approach in the design of either ssODNs, dsDNA donor cargos or both when keeping inserts in-frame with a native gene is of concern. In this study, we address this difference by implementing the necessary changes in the ssODN design and keep the dsDNA donor plasmids similar. For information on how single pair N-terminal tagging and C-terminal constructs are kept in-frame see the “dsDNA donor design” section. To keep double pair N-terminal tagging constructs in-frame, a DNA spacer is included after the second attP of the ssODN in order to influence the length of the resulting attL. The preferred sequence used in this study is the final 5 bp of the 58 bp-long attP - effectively the ssODN contains a single 48 bp attP followed by another 53 bp attP. However, an alternative spacer can be used as long as it fulfils the requirements outlined in the “dsDNA donor design” section. To keep double pair C-terminal tagging constructs inframe, a DNA spacer is included before the first attP of the ssODN in order to influence the length of the resulting attR. The sequence used in this study is the first 5 bp of the 58 bp attP variant. This effectively results in a ssODN that contains a single attP of 53 bp, followed by another 48 bp attP. However, an alternative spacer can be used as long as it fulfils the requirements outlined in the “dsDNA donor design” section. sgRNA sequences for CRISPR cuts are designed to cut as close to the insertion site as possible, whilst minimising off-target effects, and ideally with the sgRNA sequence spanning the insertion site to block recutting after correct integration. If this is not possible, synonymous or non-coding mutations should be introduced in the PAM site within the homology arms of the ssODN. sgRNAs are synthesised as chemically modified RNAs (Synthego) to minimise toxicity and maximise editing efficiency. All sgRNA sequences used in editing experiments are listed in Supplementary Table S2, and ssODN sequences used for single attP and dual attP tagging as well as BFP targeting by HDR are listed in Supplementary Table S3. For a worked example of a full tagging design see Supplementary Table S6 and Supplementary Material 1.

### dsDNA donor design

Recombination donor constructs used in this study are circular dsDNA molecules and contain attB sites which are 46 bp long (plasmid maps used in this study are listed in Supplementary Table S4). In the case of the single attP-attB pair system, a circular plasmid is cut and self-ligated to create a circular donor (containing a single attB site) - the entire length of the circularised cargo is integrated into the genome (the circularised cargo does not contain any bacterial vector backbone sequence). The parent attB cargo vector also contains single attB site and inward facing Type IIS (BbsI) restriction sites that flank the attB-insert sequence. The double attP-attB pair system contains two attB sites flanking the insert. It can be used without in vitro circularisation as upon completion of the recombination the insert cassette is inserted and the bacterial backbone sequence is excised. N-terminal tagging constructs are designed to be in-frame with the gene-of-interest post integration. The length of the attR and attL sites that result from recombination is 52 bp in the case of the single attP-attB pair system and 47 bp in the case of the dual attP-attB system. To keep N-terminal tagging constructs in-frame a DNA spacer is included in the 3’ end of the dsDNA donor construct. Importantly, the spacer shouldn’t contain stop or start codons. Sequence maps with annotated features of the tagging experiment designs for targeting exon 1 of *ACTR10* for an N-terminal mNeonGreen fusion are provided in Supplementary Table S6. and its length should complement the attL to a length divisible by 3. A quick way to check that this is possible is to perform a modulo operation with a divisor of 3 on the sequence length - it should return a remainder of 2. This is because 52 (the length of attR and attL resulting from recombination of the single attP-attB system) has a remainder of 1 when divided by 3, their sum is 54, which is divisible by 3 and hence would be in-frame in the absence of stop codons. To keep C-terminal tagging constructs in-frame a DNA spacer is included in the 3’ end of the dsDNA donor construct. As in the case of N-terminal tagging, the spacer’s length is designed to return a remainder of 2 when divided by 3 and to avoid stop codons. For a worked example of a tagging design see Supplementary Table S6.

### dsDNA donor circularisation

The dsDNA donors used in the single attP-attB pair system were generated from producer plasmids, in turn purified from bacterial cultures. In initial experiments, the plasmids were digested with BbsI and the fragment corresponding to the insert was gel purified. In subsequent experiments a PCR step to amplify the insert was performed and gel purification was omitted. Briefly, 16x 20 µL reactions were set up with 1x KAPA HS HiFi mastermix and 500 pg template plasmid DNA and primers M13F[17-mer] and IG161 at final concentrations of 300 nM each. PCR was performed with the following steps: 95°C 3 min; 25x (98°C 20 sec, 48°C 15 sec, 72°C 30 sec); 72°C 2 min. The reactions were then pooled, loaded onto a single column and purified using Macherey Nagel NucleoSpin PCR and Gel Extraction purification columns (referred to as MN kit), following the manufacturer’s protocol (in all cases, an extra step of centrifugation at 50 x g for 1 min was included during elution prior to the final spin as per manufacturer’s recommendations for increased sample recovery), elution volumes were 100-110 µL. The typical yields were between 8 and 10 µg of DNA. The pooled purified sample was digested in 5 x 100 µL reactions with 1x rCutsmart buffer (cat. #B6004) using 40 U of BbsI-HF (R0539) at 37°C overnight. The reactions were then pooled, loaded onto a single column and purified using the MN kit as above. The purified and digested insert fragment was incubated in a ligation reaction to induce circularisation. Briefly, 10x reactions of 120 µL were prepared with 1x T4 DNA Ligase Reaction Buffer (cat. #B0202) and 2400 CELU of T4 DNA Ligase and final DNA concentrations ranged between 8-9 ng/uL. The Low molar concentrations of DNA substrate and high concentrations of ligase improve the yield of the self-ligated circularised monomer. High DNA concentrations increase the accumulation of intermolecular ligation and the formation of linear and circular dimers, trimers and higher multimers. The reactions were incubated for a minimum of 2 hours up to overnight at 20°C. The reactions were then pooled, loaded onto a single column and purified using the MN kit as above. In some experiments, the ligation reactions were treated with T5 exonuclease (cat. #M0663) to remove carryover linear monomers and ligated linear multimers as well as open circles (nicked DNA) and ssODN and purified again as described above. Routine quality control of the fraction of circular monomer was performed using Agilent Tapestation D500 screentape and D5000 dsDNA kit. Initial experiments with gel-purified circular monomer resistant to T5 exonucleolytic digestion were performed on 1x TAE 1% agarose gels with EtBr to establish the correspondence of the migration pattern of the circular monomer molecular species between agarose and screentape - in the presence of EtBr, the circular monomer migrates faster than linear monomer like supercoiled DNA on agarose, on screen tapes the circular dsDNA migrates slower than linear dsDNA. For the tagging experiments shown in this paper, the molar proportion of circular monomers in the ligation end-product mixture ranged from ∼40% to 50%.

### Endotoxin removal from DNA preparations

DNA extractions were purified using the TXS method (37) prior to nucleofection. Briefly, a 0.25 volume of TXS solution was added to 1 volume of DNA solution and mixed thoroughly by inverting. The solution was then incubated at room temperature for 5-10 minutes or for as long as overnight at 4°C. A 0.25 volume of 5 M NaCl was then added and mixed thoroughly by inverting. The sample was then centrifuged for at least 10 minutes at maximum speed (>15,000 x g) at 4°C. The clear upper layer, which contains the purified plasmid, was aspirated into a clean tube (care was taken not to take up the red tinted solution at the bottom of the tube). The clear supernatant was then precipitated using 100% isopropanol, washed with 70% v/v ethanol and dried. The DNA pellet was then resuspended with nuclease-free water or TE buffer (pH 8).

### Mammalian cell culture

Kolf_2_C1 is an edited human induced pluripotent stem cell (hiPSC) line, corrected for a 19 bp deletion in one copy of ARID2, which has undergone extensive characterisation and is commonly used for editing. Kolf_2_C1 is derived by the Human Induced Pluripotent Stem Cell Initiative (HipSci) consortium. Kolf_2_C1 BFP/GFP reporter line contains a BFP reporter inserted in the ROSA26 locus under a EF1a promoter (Bassett’s lab).

The corrected A1ATD line (BOB, https://hpscreg.eu/cell-line/CAMi014-A) and Kolf_2_C1 (https://hpscreg.eu/cell-line/WTSIi018-B-1) were male hiPSC lines generated as part of Cambridgeshire 1 NRES REC Reference 09/H0304/77, Hertfordshire NRES REC Reference 08/H0311/201, London Fulham REC Reference 14/LO/0345 and 15/LO/1126), HMDMC 14/013.

hiPSC lines were cultured under feeder free condition in Stemflex medium (combo kit, Gibco TM A3349401) on Vitronectin substrate. Vitronectin was used at 1:100 dilution of a 1 mg/mL stock solution in PBS, using 1 mL per 6-well plate for coating. Incubate at RT for 1 hour before aspirating and replacing with culture media immediately. After initial thaw, cells were clump-passaged 1:10 every 4-5 days and cultured in a humidified incubator at 37°C and 5% CO2. K562 is a human erythroleukemic cell line, derived from a patient with chronic myeloid leukemia (CML), and is widely used in biomedical research. K562 cells were cultured in RPMI medium supplemented with 10% (vol/vol) fetal bovine serum (FBS).

### Nucleofection of hiPSCs and K562 cells

For genome engineering in human induced pluripotent stem cells (hiPSCs), 2×10^5^ cells were nucleofected with eSpCas9 (24 pmol) and synthetic sgRNA (Synthego, 45 pmol), an ssODN encoding the *attP* site (Ultramers from IDT, 100 pmol unless otherwise specified), a Bxb1 expression plasmid (500 ng, equivalent to 127.4 fmol unless otherwise specified), and a circular donor plasmid containing the *attB* site (variable amounts specified for each experiment). Nucleofection was performed using the P3 Primary Cell solution and program CA-137 in 16-well cuvettes on a 4D-Nucleofector X unit (Lonza) in 20 µL volumes. Cas9 and sgRNA were pre-complexed into RNPs in a 1:2 ratio - sgRNAs were diluted in IDT duplex buffer to a final concentration of 45 µM (from 200 µM stock solutions in TE buffer, pH 8.0) and eSpCas9 protein (which was produced in-house) was diluted in PBS to a final concentration of 4 mg/mL (24 µM) prior to complexing; equivalent results were achieved using commercially available high-fidelity Cas9 proteins (e.g., Cas9 HiFi, IDT).

For editing in K562 cells, the same set of reagents and concentrations was used. However, nucleofections were conducted using the SF Cell Line buffer and program FF-120, also in 16-well cuvettes using the Amaxa 4D-Nucleofector system (Lonza).

### Small molecule titration

All small molecule inhibitors tested are commercially available. AZD-7648 (MedChem Express, cat. #HY-111783), M3814 (MedChem Express, cat. #HY-101570) and NU7441 (Selleckchem, cat. #S2638) were dissolved in DMSO (Thermo Fisher Scientific) at a concentration of 5 mM. IDT HDR Enhancer v2 (IDT, cat. #10007910) was purchased as a 0.69 mM concentrated solution in DMSO. IDT HDR Enhancer v1 is no longer commercially available. Optimal concentrations of small molecule inhibitors were determined using the BFP-GFP reporter assay. Nucleofection of sgRNA (Synthego), Cas9 protein (produced in-house or HiFi Cas9, IDT) and ssODN (Ultramer DNA oligo, IDT) into the hiPSC BFP reporter line were carried out in 100 µL cuvettes (Lonza) using an Amaxa 4D-Nucleofector (Lonza), P3 Primary Cell buffer and program CA137. Final amounts per nucleofection: 1×10^6^ cells in 100 μL P3 solution, 20 μg Cas9 protein, 20 μg sgRNA and 500 pmol ssODN. Post nucleofection, cells were maintained in small molecule inhibitor supplemented culture media at various concentrations (0, 0.5, 1, 2, 4, 10, 20, 30 and 50 µM) for 24 hours. 3 days post-nucleofection, cells were analysed for presence or absence of GFP and BFP respectively by flow cytometry (CytoFLEX, Beckman Coulter).

### Genomic DNA extraction and characterisation by PCR

DNA was extracted from nucleofected cells using the DNeasy Blood and Tissue kit (Qiagen, cat. #69504) according to the manufacturer’s instructions. After purification, genomic DNA was eluted in 50 μL of water. To confirm correct integration, target regions were PCR amplified (primers in Supplementary Table S5) and analysed by gel electrophoresis. 10 ng of gDNA was used to set up 50 µL PCR reactions using KAPA HiFi HotStart ReadyMix (2x) (KAPA Biosystems, cat. #KK2601). PCR was performed with the following steps: 95°C 3 min; 35x (98°C 20 sec, 65°C 15 sec, 72°C 15 sec); 72°C 1 min.

### Genome-editing characterisation by ICE analysis for quantification of ssODN integration

A few days post-electroporation, edited cells were pelleted by centrifugation and the DNA isolated using PureLink™ Genomic DNA Mini Kit according to manufacturer’s instructions. Amplicons were generated using indicated PCR primers (Table S5), designed to amplify an approximately 1000 bp region of genomic DNA surrounding the target site. 10 ng of gDNA was used to set up 50 µL PCR reactions using KAPA HiFi HotStart ReadyMix (2x) (KAPA Biosystems, cat. #KK2601). PCR was performed with the following steps: 95°C 3 min; 35x (98°C 20 sec, 65°C 15 sec, 72°C 15 sec); 72°C 1 min. Resulting PCR product was purified using Monarch® PCR & DNA Cleanup Kit (5 μg) according to manufacturer’s instructions and shipped to Genewiz (Leipzig, Germany) for Sanger sequencing. Trace sequencing files were uploaded to ICE v2 (https://ice.synthego.com/, Synthego) to quantify efficiency of ssODN integration.

### Genome-editing characterisation by miSeq analysis

High throughput sequencing of amplicons spanning the CRISPR target sites was performed by PCR amplification of genomic DNA using methods as described in (*38*). Illumina adaptors and indexes were added in a second round of PCR followed by pooling and high throughput sequencing on a MiSeq instrument. Analysis was performed with CRISPResso2(39).

### eePassige

pegRNAs for eePassige were designed based on the method previously described (Anzalone et al. 2022 and Pandey et al. 2024). In brief, two pegRNAs were designed on opposing strands of the target locus with a view to deleting the short intervening sequence while at the same time installing an attP recombination site. All reagents required for the eePassige experiment were delivered by nucleofection. The enzymatic components were delivered as mRNA, the two pegRNAs as synthetic RNAs supplied by IDT and the Bxb1 recombinase as an expression plasmid. The DNA cargo constructs for recombination with the attP site once inserted into the genomic locus were either plasmid or minicircle (details in figure legends)

### HDR with long ssODN template

The template for ssODN production was prepared by PCR amplifying the fragment, containing homology arms from plasmid template pCR+mNeonGreen-ACTR10 with primers ACTR10-Neon_F3 (5’ phosphorylated) and ACTR10-Neon_R3. The reactions were pooled, digested with DpnI and cleaned using a Zymo DNA Clean & Concentrate kit. The purified PCR template was treated with Takara long ssODN Strandase kit according to the manufacturer’s instructions. The reactions were pooled and cleaned up with Zymo DNA Clean & Concentrate kit and then precipitated with EtOH, washed and resuspended. The ssODN used was 1185 nt long and was resuspended in a small volume (typical yield - 5 µg ssODN in 3 µL TE buffer, pH 8.0). The full 5 µg of DNA was used to transfect 2×10^5^ iPS cells as per iPSC nucleofection method described above, omitting ssODN, Bxb1 expression plasmid and a circular donor plasmid.

### ddPCR

5 to 20 ng of genomic DNA was used per 20 µL reaction including 10 μL of ddPCR Supermix for probes (no dUTP) (Bio-Rad, cat. #186-3023) and 0.5 μL of VIC-tagged TaqMan copy number reference assay, human RNase P (Thermo Fisher Scientific, cat. #4403326) as a reference. 2 pmol each of forward and reverse primers as well as 4 pmol of a probe for the target locus were used per 20 µL reaction. All the primers and probes used for ddPCR are listed in Supplementary Table S5. Droplets were generated using a QX200 Droplet Generator (Bio-Rad, cat. #1854002) and transferred to 96 well plates. PCR was performed with the following steps: 95°C 3 min; 40x (95°C 20 sec, 60°C 1 min); 98°C 10 min.

The resulting FAM and VIC fluorescence was read using a QX200 Droplet Reader (Bio-Rad, cat. #1864003) and positive droplets were determined. The copy number of the target locus was calculated as the ratio of the positive droplets for the target locus to those of the reference gene, multiplied by 2 (copy number of the reference gene).

### Nanopore sequencing using Cas9 enrichment

Kolf 2.1s cells were targeted with a mNeonGreen construct at the *ACTR10* locus using either the single attP (GT) cassette or the dual attP (GA-GT) cassettes. Cells were then sorted on green fluorescence after a week of growth, following transfection. Five million cells per condition (sorted and unsorted, single and dual-cassette) were pelleted and frozen at −20°C. The pellets were thawed and high molecular weight (HMW) DNA was extracted using the Qiagen Magattract HMW kit (cat. #67563), following the manufacturer’s protocol for blood cells. The HMW DNA was checked for quality and integrity on a Tapestation 4150 using a Genomic DNA screentape. The DNA extractions were found to >60 kb in length. The HMW DNA was then processed in accordance with the Oxford Nanopore Technologies protocol ENR_9084_v109_revW_04Dec2018. The sgRNAs used to cut outside of the *ACTR10* locus are listed in Supplementary Table S2. Analysis of sequencing data was performed using Knock-Knock software (40).

### Imaging

Cells for imaging were plated on PhenoPlate 96-well black walled microplates (Revvity, cat. #6005182). Once confluent cells were washed with 100 µL PBS per well and fixed at room temperature for 10 minutes with 4% PFA supplemented with Hoechst at 1 µg/mL for nuclei staining (40 µL/well). Post-fixing, cells were washed 3 times then stored in 100 µL PBS. Cells were imaged on an Opera Phenix (Perkin Elmer) using a 40x/1.1 NA water lens, or as specified in the figure legends. Hoechst and mNeonGreen were excited at 375 nm and 488 nm, respectively. Hoechst fluorescence was collected at 435-480 nm with an exposure time of 60 ms, and mNeonGreen fluorescence was collected at 500-525 nm with an exposure time of 100 ms.

### Primary T cell isolation and stimulation

Human biological samples were sourced ethically, and their research use was in accord with the terms of informed consent under an institutional review board/ethics committee-approved protocol (15/NW/0282). Peripheral blood mononuclear cells (PBMCs) were isolated from fresh leukapheresis products from human healthy donors (Leukopaks, BioIVT) using a Ficoll-Paque PLUS (GE Healthcare, cat. #GE17-1440-03) density gradient centrifugation. Cells were cryopreserved in freezing media (RPMI 1640 (Gibco, cat. #52400025), 10% DMSO, 50% Fetal Bovine Serum (FBS, SIGMA-ALDRICH, cat. #F9665)) and stored in liquid nitrogen. PBMCs were thawed a day before T cell isolation, resuspended in complete RPMI media at 20e6 cells/mL (RPMI 1640, 10% FBS, 100 U/mL Penicillin-Streptomycin (Gibco, cat. #15140122), 2mM L-Glutamine (Merck, cat. #G7513)) and incubated at 37°C 5% CO2 overnight. PBMCs were collected and washed twice with DPBS without calcium or magnesium (Gibco, cat. #14190144). Memory CD4+ T cells were isolated by immunomagnetic negative selection using the EasySep™ Human Memory CD4+ T Cell Enrichment Kit (STEMCELL Technologies, cat. #19157) according to the manufacturer’s instructions. After isolation, cells were cultured in media consisting of StemPro™-34 SFM (Gibco, cat. #10639011), 10% FBS, 100 U/mL Penicillin-Streptomycin, 2mM L-Glutamine, recombinant Human IL-2 at 15 ng/mL (R&D systems, cat. #10453-IL) (named as complete StemPro) at 1e6 cells/mL. Cells were then stimulated with Dynabeads® Human T-Activator CD3/CD28 at a 1:1 bead:cell ratio (Thermo Fisher Scientific, cat. #11161D).

### Nucleofection of primary T cells

Lyophilized sgRNAs (Synthego) and ssODNs (IDT) were resuspended in water to a stock concentration of 100 µM and stored at −20°C until use. RNPs were produced by mixing sgRNAs (180 pmol per reaction) and Cas9 (Alt-R™ S.p. Cas9 Nuclease V3, IDT, 61 pmol per reaction) at a 3:1 sgRNA:Cas9 molar ratio. The following ONE-STEP components were added to the RNP complexes: ssODN GA GT donor template (100 pmol), eeBxb1 plasmid (500 ng), and EGFP TRAC plasmid (2.1 µg). Stimulation beads were removed by placing primary T cells on a magnet for 1 min and collecting the supernatant. Cells were then spun down for 5 min at 400 x g and washed twice with DPBS without calcium or magnesium, before being resuspended in B1mix buffer (buffer B1 (5 mM KCl, 15 mM MgCl2, 120 mM Na2HPO4/NaH2PO4 pH 7.2, 10 mM sodium succinate and 25mM mannitol) were combined with mix solution (0.5 mM sodium pyruvate, 0.8 mM Ca(NO3)2, 0.26 mM i-inositol (myo-inositol), 4 mM GlutaMAX (L-alanyl-L-glutamine dipeptide in 0.85% NaCl), 20 mM d-glucose) in a v/v ratio of 72.3 to 27.7) at 7e5 cells per 20 µL and added to the ONE-STEP pre-mix. The cells in the buffer were then transferred to a 16-well Nucleocuvette™ Strip (Lonza, 4D-Nucleofector™ X Kit) for nucleofection using the pulse code EH-115. Immediately after nucleofection, 80 µL of pre-warmed complete StemPro media was added to each well and incubated at 37°C with 5% CO2 for 15 min. The cells were then transferred to a 96-well round-bottom plate containing 145 µL of media with 0.5 µM AZD-7648 and incubated at 37°C with 5% CO2 for 24 hours. AZD-7648 was then washed out by removing supernatants without disturbing the pellets and resuspending the cells in 250 µL of pre-warmed media.

## Results

### 1. ONE-STEP tagging combines CRISPR-Cas9 editing with Bxb1 site specific integration

We have devised a versatile integration system that combines the precision of CRISPR-Cas9-based HDR with the integration of large DNA cargos via the Bxb1 serine integrase. Bxb1 is functional in mammalian cells and efficiently catalyzes unidirectional (non-reversible) integration between double-stranded DNA (dsDNA) sequences containing an attP and their complementary attB attachment site. By utilising CRISPR-Cas9 to position the integrase attP sites at specific genomic locations, we can direct Bxb1 (delivered in *trans*) to act at the chosen sites. By simultaneously providing a circular double-stranded DNA template containing the attB attachment site, we aim to achieve direct integration in a single step (Fig. 1: schematic).

**Figure 1:**
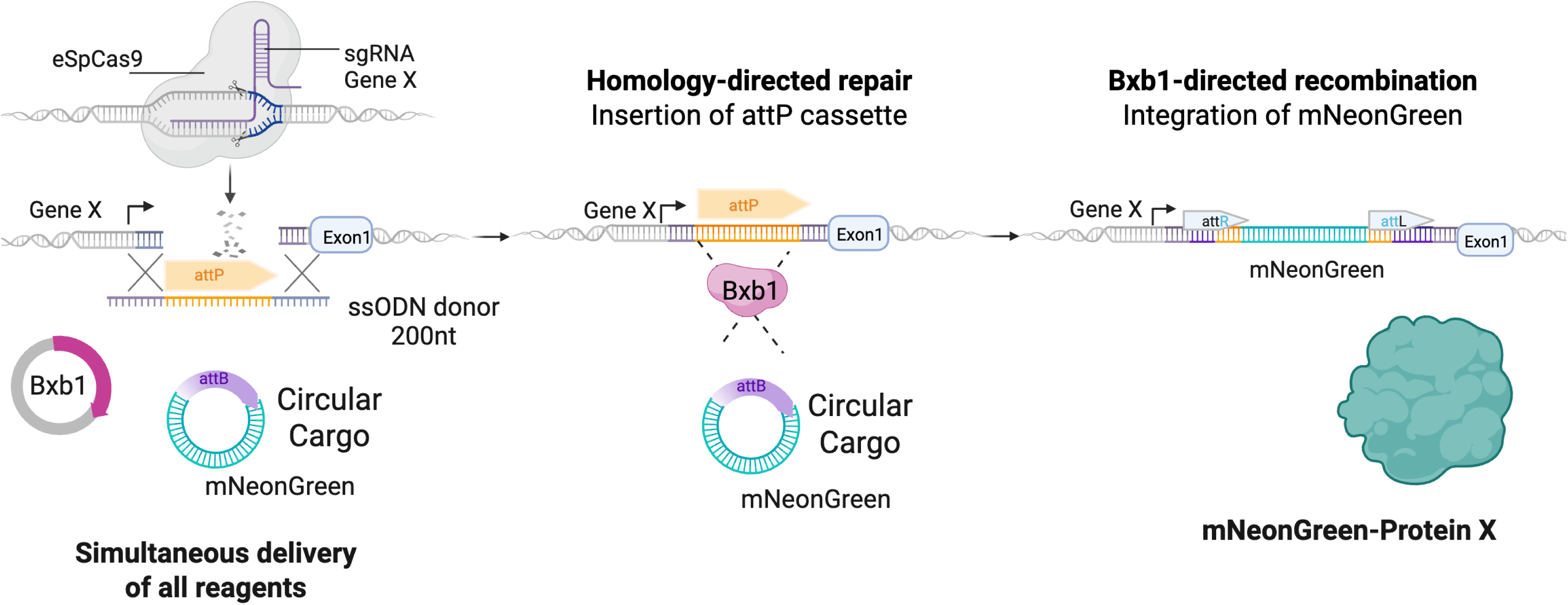
Schematic of the ONE-STEP tagging technology. The system involves insertion of landing site via Crispr-Cas9 mediated by homology directed repair (HDR) and a single stranded DNA oligo (ssODN) as template. Upon successful HDR, the Bxb1 integrase results in site-specific integration of the mNeonGreen cargo. All reagents are delivered simultaneously.

Because Bxb1 specifically recognizes dsDNA, it cannot recombine attP sites delivered as single-stranded oligonucleotides (ssODNs) until they have been integrated into the genome and converted into dsDNA. As a result, recombination with the dsDNA cargo donor only occurs after successful integration of the attP site. Following recombination, Bxb1 leaves behind short residual sequences—attL and attR—at the genomic integration site and we additionally include a flexible (GGSGGGSG) linker to isolate the tag from the gene of interest. In our experiments, the attB-containing cargo was delivered on a minicircle vector.

To validate our method, we designed an experiment to endogenously tag the N-terminus of the *ACTR10* gene with a fluorescent protein encoding gene (mNeonGreen) in human induced pluripotent stem cells (hiPSCs). All components required for integration were delivered simultaneously in a single nucleofection. These included a CRISPR-Cas9 ribonucleoprotein complex targeting *ACTR10*, a short single-stranded HDR template (200 nt) containing an attP site flanked by ∼70 bp homology arms, a plasmid encoding Bxb1 recombinase, and a circular dsDNA cargo bearing an attB site and the mNeonGreen sequence. To avoid plasmid backbone integration, the donor cargo was prepared as a minicircle via self-circularisation of a restriction fragment.

Seven days post-nucleofection, we evaluated integration outcomes by flow cytometry and PCR-based genotyping, with each condition tested in triplicate. Titrating the circular donor cargo at increasing molar excesses (1x, 3x, and 6x relative to the Bxb1 plasmid) yielded a dose-dependent increase in tagging efficiency, reaching approximately 10% at the highest dose. In contrast, omitting the attP-containing ssODN HDR template resulted in minimal random integration (∼0.1%) even at the highest cargo concentration, confirming the specificity of the recombination reaction (Fig. 2a). Successful integration was validated by PCR (Fig. S1a), Sanger sequencing (see alignments in the “Integrated mNeonGreen *ACTR10* Design” map listed in Supplementary Table S4 and Supplementary plasmid maps) and long read Nanopore sequencing using Cas9 enrichment (Fig. S1b). This showed that the majority of events after sorting were as expected with a precise integration of the mNeonGreen cassette but with a low frequency of indels at the integration junctions.

**Figure 2:**
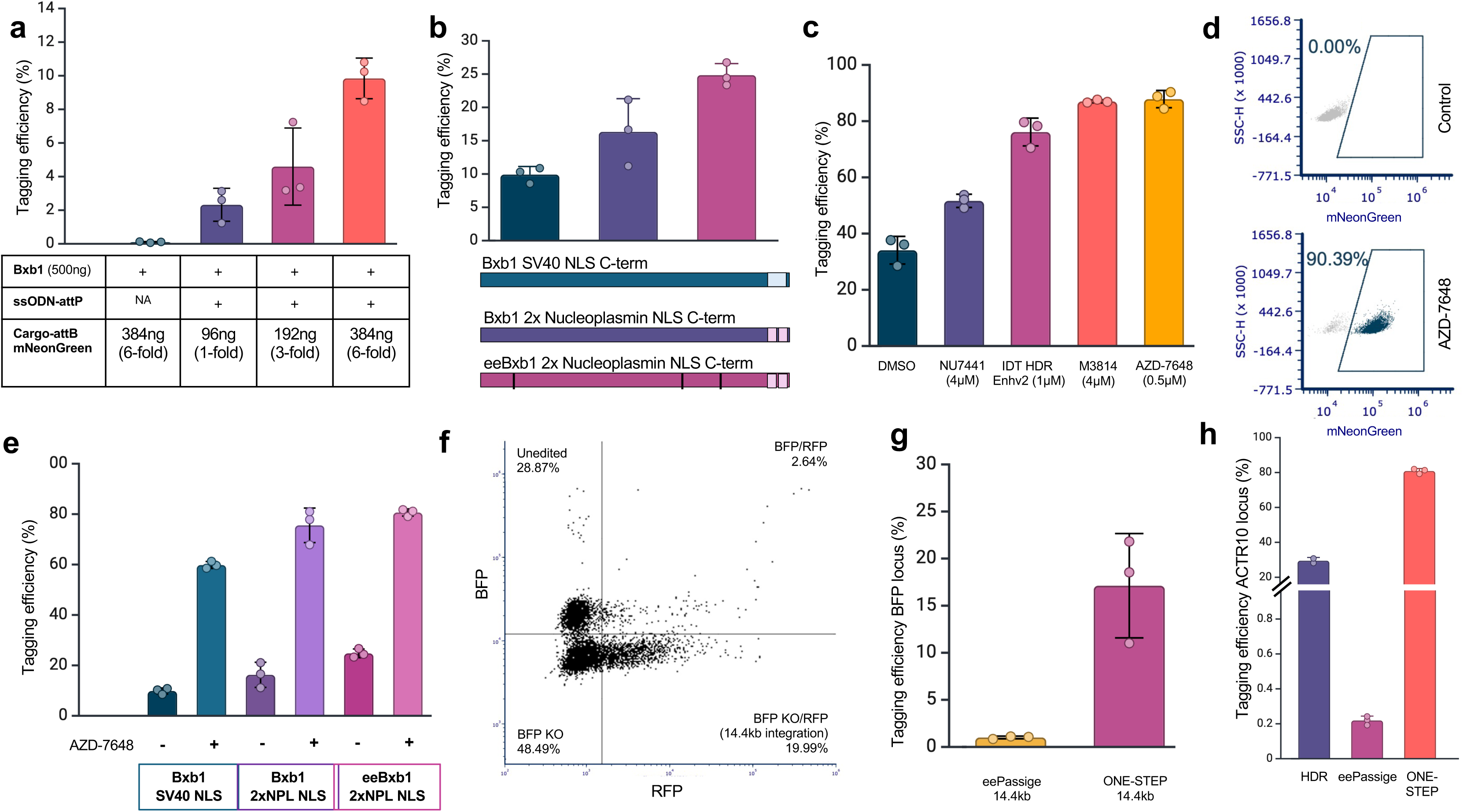
Optimisation of ONE-STEP tagging for efficient cargo integration. **a.** Tagging efficiency at the *ACTR10* locus using different concentrations of circularised mNeonGreen donor plasmid (1-, 3-, or 6-fold molar excess relative to Bxb1), with a constant 500 ng of Bxb1 plasmid. The “Tagging efficiency” condition includes all required components; the “Random integration control” omits the ssODN-attP (NA), resulting in <1% non-specific integration. A 6-fold excess of mNeonGreen donor achieved ∼10% tagging efficiency (n=3). Error bars represent standard deviation. **b.** Comparison of tagging efficiency at the *ACTR10* locus using different Bxb1 variants and nuclear localisation signals (NLS): WT Bxb1 with C-terminal SV40 NLS, WT Bxb1 with C-terminal 2×NPL NLS, and engineered eeBxb1 with C-terminal 2×NPL NLS. The eeBxb1 2×NPL NLS variant achieved ∼25% tagging efficiency with a 6-fold excess of donor (n=3). Error bars represent standard deviation. **c.** Effect of DNA-PK inhibitors on tagging efficiency. Cells were treated immediately after nucleofection, with media replacement after 24 hours. Compounds tested: NU7441 (4 μM), M3814 (4 μM), AZD-7648 (0.5 μM), and IDT HDR Enhancer v2 (1 μM). AZD-7648 (0.5 μM) treatment resulted in >80% tagging efficiency at the *ACTR10* locus (n=3). Error bars represent standard deviation. **d.** Representative flow cytometry plots showing *ACTR10*-mNeonGreen tagging efficiency in untreated control cells (top) and cells treated with AZD-7648 (bottom). **e.** Tagging efficiency at the *ACTR10* locus using a 6-fold molar excess of mNeonGreen donor with different Bxb1 variants/NLS combinations,with or without AZD-7648. The effect of AZD-7648 was variant-dependent, with the largest enhancement observed for WT Bxb1 with SV40 NLS. **f.** Bar chart showing integration of a 14.4 kb cargo into the BFP reporter gene integrated as a single copy at the ROSA26 locus in the Kolf_2_C1 hiPSC line. ONE-STEP tagging achieved ∼16% efficiency, compared to ∼1% using eePASSIGE (n=3). Error bars represent standard deviation. **g.** Representative flow cytometry plot illustrating 14.4 kb integration at the BPF reporter gene. Successfully tagged cells exhibited loss of BFP and gain of RFP signal. h. Benchmarking of ONE-STEP tagging against HDR and eePASSIGE at the *ACTR10* locus. mNeonGreen integration efficiency was ∼30% with HDR (n=2), ∼0.2% with eePASSIGE (n=3), and ∼80% using ONE-STEP tagging (n=3). Error bars represent standard deviation.

These findings demonstrate that efficient, site-specific tagging of *ACTR10* with mNeonGreen can be achieved through a streamlined, single-step delivery of all necessary components. Based on this, we refer to this approach as ONE-STEP tagging. Moreover, our results show that increasing the concentration of the circular cargo alone significantly improves integration efficiency, without compromising specificity (Fig. 2a).

### 2. Enhancing tagging efficiency by improving nuclear localisation and catalytic activity of Bxb1

We next addressed whether improving the nuclear localization of Bxb1 could enhance its integration efficiency, similar to the optimization efforts previously undertaken for CRISPR-Cas9 which showed that the choice and combination of nuclear localization signals (NLS) significantly influenced its efficiency (41–43). Our original wild-type (WT) Bxb1 plasmid includes a SV40 C-terminal NLS, but both our lab and others (44) have observed that adding two bipartite nucleoplasmin (NPL) NLS at the C-terminus substantially increases efficiency, nearly doubling it in the specific case of *ACTR10* N-terminal tagging. This enhancement is even more pronounced when the C-terminal 2xNPL NLS is fused to eeBxb1, the most catalytically active variant of Bxb1 (30), resulting in tagging efficiencies of up to 25% (Fig. 2b). Based on these findings, we will consistently utilize 2xNPL eeBxb1 and a 6-fold molar excess of donor cargo in future experiments to maximize integration efficiency.

### 3. AZD-7648 is a potent DNA-PK inhibitor that improves gene tagging in hiPSCs

NHEJ and HDR are the two primary DNA damage response pathways responsible for processing DSBs. NHEJ, characterized by the modification and ligation of blunt DNA ends, functions throughout the cell cycle and operates independently of sequence homology. It is kinetically faster than HDR-related mechanisms and serves as the predominant DSB repair pathway in many cell types. In contrast, HDR is restricted to the S- and G2-phases of the cell cycle, where the presence of a sister chromatid as a template enables accurate and potentially error-free repair.

Efforts to enhance template-based repair pathways have focused on regulating key DNA repair factors, modulating CRISPR-Cas9 components, and altering the intracellular environment at DSB sites (Charpentier et al., 2018; Rees et al., 2019; Aird et al., 2018). NHEJ suppression through knockout of *lig4* was shown to improve HDR rates in *D. melanogaster* (*45*). Inhibiting DNA-PK, a crucial factor in the NHEJ pathway, is particularly effective at enhancing CRISPR-mediated insertions (46, 47). Based on these findings, we evaluated the effects of four commercially available DNA-PK inhibitors (NU7441, AZD-7648, M3814, and IDT HDR Enhancer v2) to promote HDR-mediated repair and thus improve tagging efficiency.

We first optimized the concentrations of DNA-PK inhibitors using a human induced pluripotent stem cell (hiPSC) reporter line (Kolf_2_C1) containing a single copy of blue fluorescent protein (BFP) integrated at the hROSA26 locus. This reporter system allows for precise targeting of the fluorophore binding site by CRISPR-Cas9 to generate a DSB. Repair via NHEJ results in indel formation and loss of BFP fluorescence. In contrast, when an HDR template introducing a two-nucleotide mutation (S66T, H67Y) is used, successful repair leads to a shift from blue to green fluorescence, providing a quantifiable measure of HDR activity (BFP-GFP reporter assay) (Fig. S2).

After optimizing the concentrations (NU7441, 4 μM; M3814, 4 μM; AZD-7648, 0.5 μM; IDT HDR Enh v2, 1 μM), we tested the effect of each of the four DNA-PK inhibitors on tagging efficiency by enhancing HDR rates, specifically by inserting the ssODN containing the attP attachment site via HDR. Among the four inhibitors tested, AZD-7648 (0.5 μM) demonstrated the most pronounced improvement in tagging efficiency and at the lowest concentration when using eeBxb1 with the 2x NPL NLS, resulting in a 3-fold increase compared to the DMSO control as shown by the bar chart (Fig. 2c) and representative flow cytometry plots (Fig. 2d). The effect of AZD-7648 was observed with all three Bxb1 variants (Fig. 2e). Treatment with the optimised concentration of AZD-7648 DNA-PK inhibitor increased tagging efficiency up to 90%. HDR rates (attP integration) of three further sites (MAP4, LMNA and FBL) were analysed by Inference of CRISPR Edits (ICE)(48), PCR genotyping and miSeq which confirmed improved rates of HDR with the use of AZD-7648 (Fig S3) although the absolute values from the different methods vary. This is likely due to the sensitivity of the different methods, especially to low frequency events which certain methods such as ICE cannot detect.

### 4. ONE-STEP tagging is less affected by cargo size

A major advantage of recombinase-based tagging is that integration efficiency is less sensitive to cargo size compared to HDR-based methods. To test the system’s capacity for large payloads, we engineered a 14.4 kb plasmid containing an attB site and multiple reporter genes. This construct included two divergent constitutive promoters: one driving a tagBFP reporter gene on one side, and the other driving both RFP and GFP on the opposite side. The attB site was strategically placed between the promoter and the tagBFP gene (Fig. S4).

We then targeted this plasmid for integration into the same hiPSC BFP reporter line. Successful site-specific integration into this locus disrupts the genomic BFP reporter as well as the tagBFP transgene on the plasmid, since the attB insertion separates the promoter from the coding sequence. In contrast, the RFP and GFP expression cassettes on the plasmid remain intact and serve as markers of successful integration. Using this assay, we observed precise integration of the full 14.4 kb construct in 16.6% of cells, as confirmed by flow cytometry and genotyping (Fig. 2f, 2g and SUPP). These results highlight the robustness of the ONE-STEP approach for delivering large genetic payloads into defined genomic loci with high specificity and efficiency. We benchmarked our system against prime editing approaches like eePassige, which resulted in only 1% integration efficiency under the same conditions (Fig. 2f). These results align with the very low integration efficiency observed for eePassige in human iPSCs, as reported by David Liu’s lab (30), where they achieved an average of only 4% efficiency when integrating a 5.6 kb donor plasmid into the *CCR5* locus of human iPSCs. In addition, we compared our ONE-STEP method with classical HDR with long ssODN template and eePassige at the *ACTR10* locus. HDR achieved 30% integration efficiency, while ONE-STEP tagging resulted in ∼80% efficiency. In contrast, eePassige yielded less than 0.5% integration efficiency (Fig. 2h). We also tested NHEJ-mediated approaches, including HITI, but were unsuccessful, with no tagging observed. These comparisons highlight the potential advantages of the ONE-STEP method in terms of integration efficiency of large constructs in human iPSCs.

### 5. Multiplexed ONE-STEP tagging

The central dinucleotide within the attP and attB sites of Bxb1 integrase plays a crucial role in the recombination process by facilitating specific pairing between these attachment sites (Ghosh et al., 2003). Modifying the central dinucleotide sequence can alter integrase activity and confer orthogonality, enabling the targeted integration of different fluorophores at distinct genomic loci (Jusiak et al., 2019). Several dinucleotide variants—such as GA, CT, and AG—have been reported to exhibit higher integration efficiencies than the wild-type GT sequence, with many achieving efficiencies greater than 75%, making them promising candidates for multiplexed integration strategies (29).

To explore this, we first assessed mNeonGreen integration efficiency at the *ACTR10* locus using different attP site variants. Specifically, we compared the wild-type attP-GT with an attP-GA variant. Our goal was to evaluate the specificity of matched versus mismatched attB/attP dinucleotide interactions (Fig. 3a). We found that both GT and GA attP sites supported similarly efficient cargo integration when paired with the corresponding attB variant. In contrast, mismatched combinations led to significantly reduced integration (∼10-fold lower), indicating minimal crosstalk between the variants (Fig. 3b).

**Figure 3.**
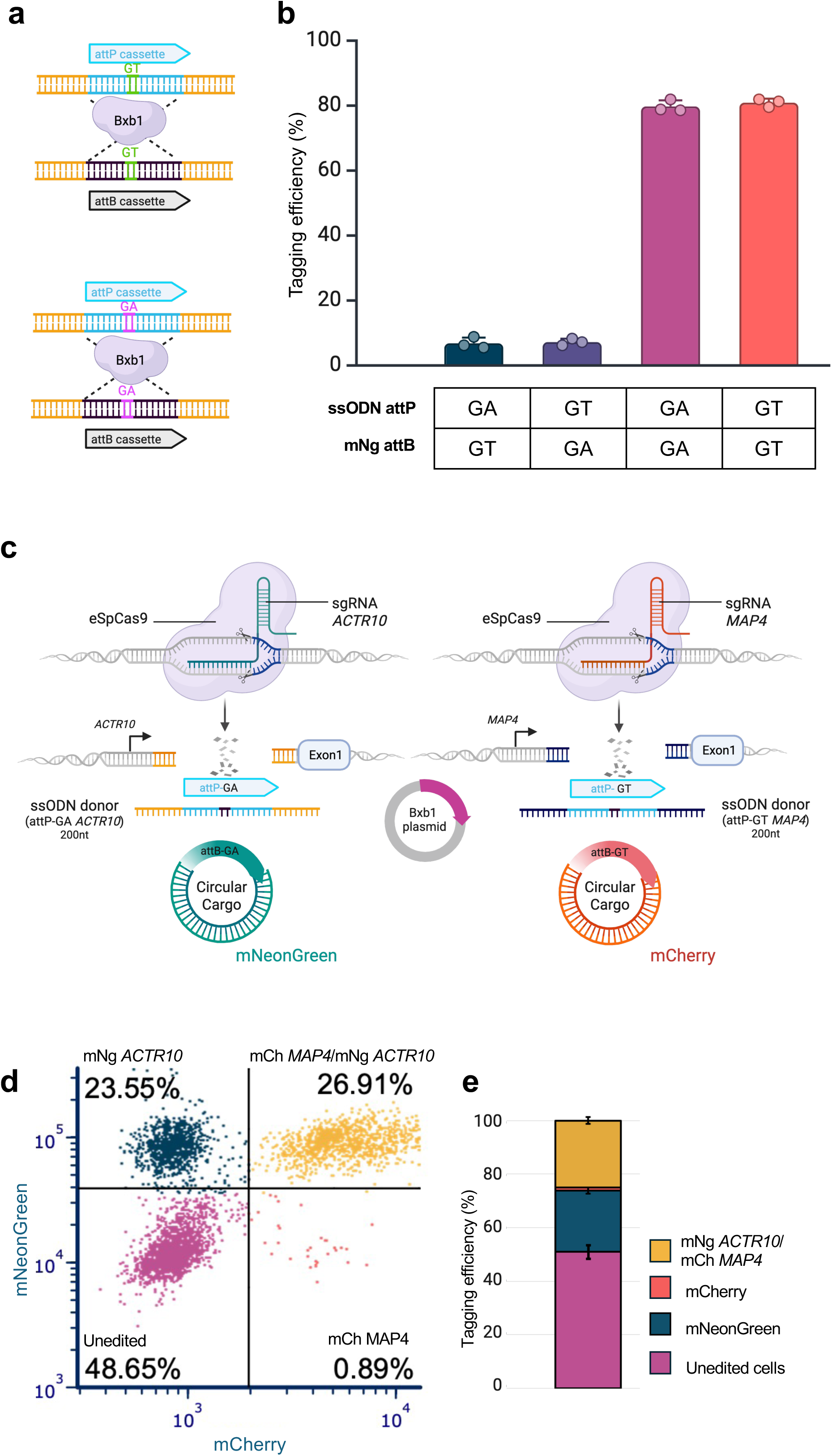
Efficient and specific integration using heterotypic recombination sites enables multiplexed tagging. **a.** Schematic representation of attP/attB-GA and attP/attB-GT cassettes. **b.** Evaluation of mNeonGreen integration at the *ACTR10* locus using attP-GA or attP-GT variants and matched or mismatched attB/attP dinucleotide combinations. Specific pairing of attB and attP dinucleotides (GA–GA or GT–GT) results in higher integration efficiency compared to mismatched pairs. **c.** Schematic of the multiplexed protein tagging strategy. **d.** Representative flow cytometry plot showing integration efficiencies at the *ACTR10* locus (mNeonGreen), *MAP4* locus (mCherry), and simultaneous integration at both loci (double-positive for mNeonGreen and mCherry). **e.** Stacked bar chart showing quantification of flow cytometry data from n=3 biological replicates. Error bars represent standard deviation. A substantial population of double-positive cells (up to ∼26%) was observed.

A key application of orthogonal integrase systems is multiplexed protein tagging, enabling simultaneous visualization of protein localization and interaction within individual cells. To demonstrate this, we used the ONE-STEP tagging method to simultaneously tag *ACTR10* with mNeonGreen and *MAP4* with mCherry in the same cell (Fig. 3c). Flow cytometry analysis confirmed successful dual tagging, with a significant population of double-positive cells (up to ∼26%, Fig. 3d, 3e).

### 6. ONE-STEP tagging and simultaneous CRISPR editing

Another valuable application of the ONE-STEP tagging technology is its ability to combine targeted protein tagging with a simultaneous CRISPR-mediated genome editing event—such as HDR or a KO—at a second locus.

To demonstrate this, we firstly delivered reagents designed to tag *MAP4* with mCherry in the BFP reporter hiPSC line described above. The presence of mCherry fluorescence confirmed successful tagging, while the BFP-to-GFP reporter system enabled simultaneous assessment of editing efficiency at a second locus. Specifically, gain of GFP signalled an HDR event (Fig. 4a). By flow cytometry, we observed efficient mCherry integration at the *MAP4* locus (∼10%), 80% of which also had HDR outcomes in the reporter assay, resulting in ∼8.5% where both events occurred simultaneously (Fig. 4b, 4c). More of the mCherry *MAP4* cells showed an HDR event (80%) than untagged cells (30%) (Fig. 4d) showing that selection for successful tagging can be used to enrich other editing events.

**Figure 4.**
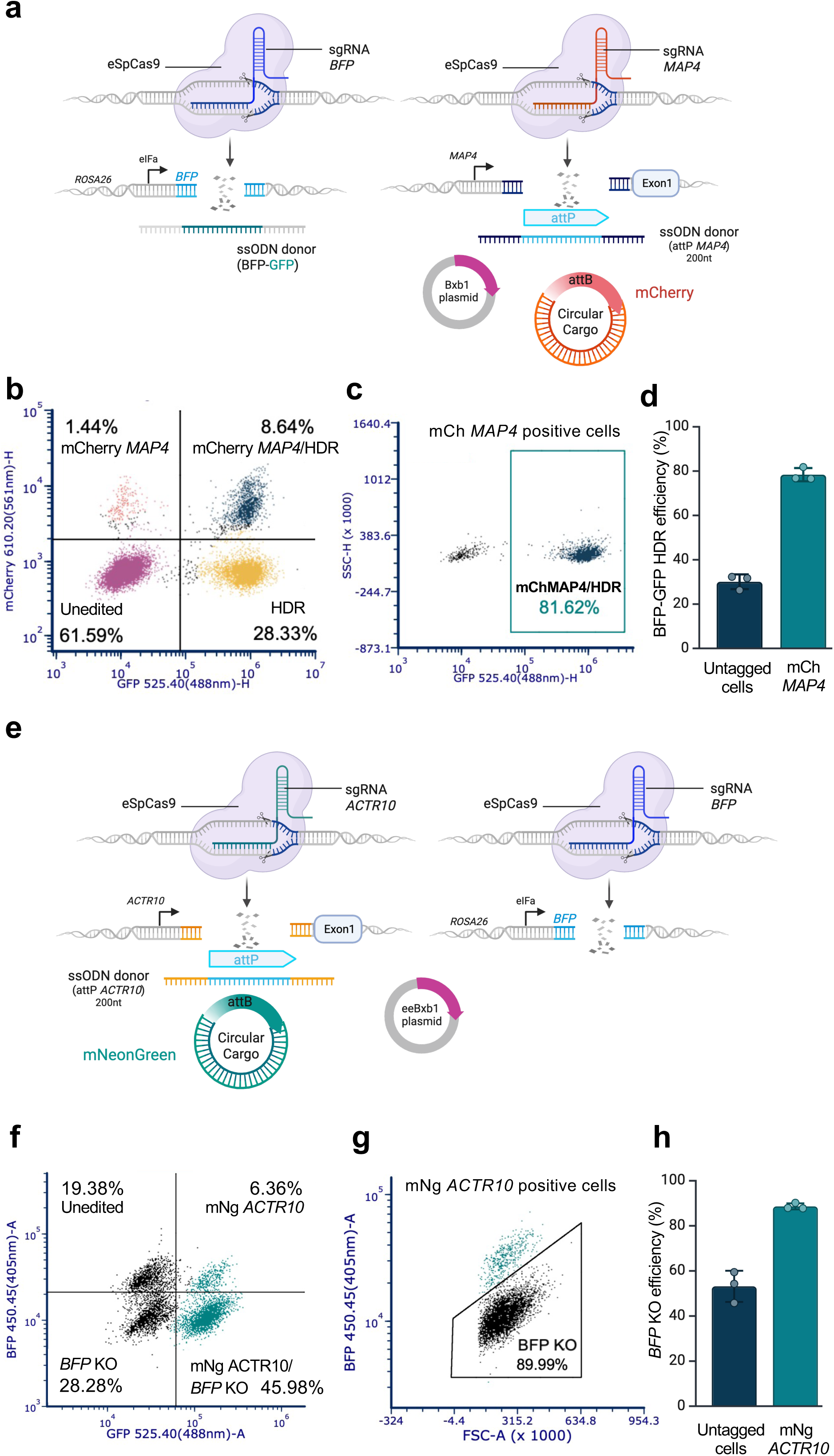
Simultaneous integration and genome editing using ONE-STEP tagging. **a.** Schematic of experimental design. Reagents for tagging *MAP4* with mCherry were co-delivered with a sgRNA and HDR template to convert BFP to GFP in a BFP reporter cell line. The mCherry signal indicates *MAP4* tagging, while GFP gain monitored HDR at the BFP locus. **b.** Representative flow cytometry plot showing integration at the *MAP4* locus (mCherry) and simultaneous HDR at the BFP locus (GFP); ∼8% double-positive cells were detected. **c.** Representative flow cytometry plot showing that ∼80% of mCherry-positive cells exhibited GFP conversion, indicating successful HDR. **d.** Bar chart quantifying HDR events in mCherry-tagged versus untagged cells. Approximately 80% of mCherry-positive cells showed HDR, compared to ∼30% of untagged cells. Data represent n=3 biological experiments; error bars indicate standard deviation. **e.** Schematic of experimental design for simultaneous tagging of *ACTR10* with mNeonGreen and knockout (KO) of the BFP gene. mNeonGreen-tagging reagents for *ACTR10* were co-delivered with a BFP sgRNA into the BFP reporter line. Successful KO events were monitored by loss of BFP expression. **f.** Representative flow cytometry plot showing integration at the *ACTR10* locus (mNeonGreen) and simultaneous BFP KO; ∼46% of cells were mNeonGreen-positive and BFP-negative. **g.** Representative flow cytometry plot showing that ∼90% of ACTR10-mNeonGreen-positive cells exhibited BFP loss. **h.** Bar chart quantifying KO events in mNeonGreen-tagged versus untagged cells. Approximately 90% of mNeonGreen-positive cells exhibited BFP loss, compared to ∼50% of untagged cells. Data represent n=3 biological experiments; error bars indicate standard deviation.

We then process to deliver reagents designed to tag *ACTR10* with mNeonGreen in the same BFP-based reporter cell line, this time looking for a simultaneous loss of BFP indicating a NHEJ-mediated knockout (Fig. 4e). Again, we observed efficient mNeonGreen integration at the *ACTR10* locus (∼52%), ∼90% of which (∼46%) had simultaneous BFP KO in the reporter assay (Fig. 4f, 4g). Again, selection for tagged cells enriched for knockout (∼90%) in comparison to the untagged population (∼53%) (Fig. 4h).

These two sets of experiments validate the ability of the ONE-STEP approach to support multiplexed genome engineering in a single experiment.

### 7. Versatile tagging at clinically relevant site in primary T cells

We decided to extend the ONE-STEP tagging system to edit more clinically relevant cells, such as primary human T cells, at the therapeutically relevant T cell receptor (TCR) locus. To achieve best efficiency of tagging, we modified nucleofection conditions using higher concentrations of Cas9, sgRNA and ssODN HDR template. The P3 buffer (Lonza) was replaced by an optimised nucleofection buffer (B1mix) which exhibits superior performance in delivering DNA to T cells (<u>An et al., 2023</u>). We successfully integrated a cargo of 4.4 kb (expressing EGFP) using a ssODN containing the attP site as HDR template, and sgRNA targeting the *TRAC* locus (Fig. 5a). We observed a tagging efficiency of ∼12% (Fig. 5b), suggesting the potential to adapt this technology for the introduction of chimeric antigen receptor (CAR transgenes at the TCR locus in primary T cells. This advancement could enhance current strategies for cancer immunotherapy through adoptive transfer.

**Figure 5.**
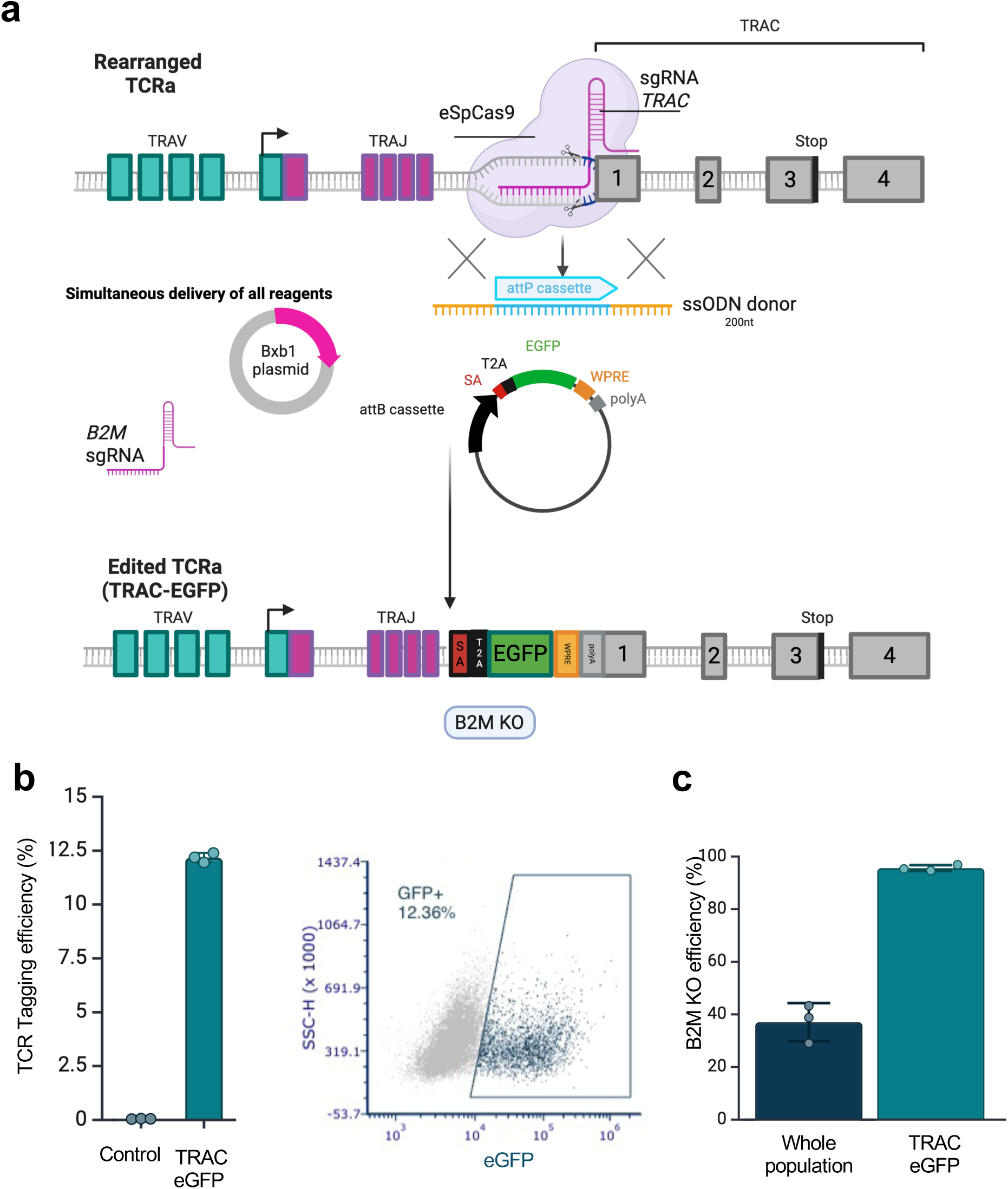
Targeted integration at the TCR locus and simultaneous knockout of B2M. **a.** Schematic of experimental design. An eGFP-expressing 4.4 kb cargo was integrated into the *TRAC* locus using a single-stranded DNA oligonucleotide (ssODN) containing an attP site as an HDR template, together with a sgRNA targeting the *TRAC* locus. A B2M-targeting guide RNA can be co-delivered to achieve simultaneous knockout at the *B2M* locus. **b.** Left: Bar chart showing ∼12% tagging efficiency at the *TRAC* locus. Data represent triplicate experiments; error bars indicate standard deviation. Right: Representative flow cytometry plot illustrating successful eGFP integration. **c.** Schematic and flow cytometry analysis of simultaneous eGFP integration at the *TRAC* locus and *B2M* knockout (KO). All components were delivered in a single nucleofection. Seven days post-nucleofection, Flowcytometry analysis revealed that >90% of eGFP-positive cells (indicating successful TRAC targeting) lacked *B2M* expression, compared to ∼30% *B2M* KO in the total cell population.

Beyond proof-of-concept applications for ONE-STEP tagging and simultaneous CRISPR editing, this dual-function system enables more sophisticated genome engineering strategies. For example, it can be used to insert a CAR construct under the control of the endogenous TCR promoter while simultaneously knocking out immune-evasive genes such as *B2M* (Fig. 5a). Disruption of *B2M* allows engineered T cells to evade recognition and destruction by alloresponsive T cells (49).

To test this approach, we delivered all components in a single nucleofection to integrate an EGFP-expressing transgene at the *TRAC* locus and simultaneously knock out *B2M*. Seven days post-nucleofection, flow cytometry analysis revealed that over 90% of EGFP-positive cells (i.e., those with successful *TRAC* targeting) also lacked *B2M* expression, compared to only ∼30% *B2M*-KO in the overall cell population (Fig. 5c). Thus, selecting for the tagging event also offers a convenient way to enrich for simultaneous knockout in the same cells.

This streamlined strategy demonstrates a powerful and scalable route for generating off-the-shelf CAR-T cells, enabling precise transgene insertion and concurrent disruption of immunogenicity-associated genes in a single step.

### 8. Dual-cassette ONE-STEP tagging using a combination of heterologous attachment sites

To increase the versatility of the technology, we developed a dual-cassette tagging strategy that eliminates the need to generate a new minicircle for each donor construct. This approach uses a ssODN HDR template (∼200 nt) that incorporates two heterologous attP sites (attP-GA and attP-GT), while maintaining ∼50 bp homology arms on either side. This universal template is used in combination with a plasmid donor in which the cargo is flanked by the corresponding variant attB sites (Fig. 6a). We tested mNeonGreen integration efficiency at the *ACTR10* locus, comparing the original ONE-STEP technology with the dual-cassette ONE-STEP tagging strategy. While the dual-cassette strategy demonstrated efficient tagging (60%), it showed a slightly reduced efficiency compared to the original method (80%) (Fig. 6b). This reduction is likely due to the lower molar concentration of the donor, as the plasmid backbone is present alongside the cargo (unlike the minicircle). We have shown (Fig. 2a) that this is critical for tagging efficiency, and we cannot further increase the donor concentration without negatively impacting cell viability (DNA toxicity). Additionally, there is likely a reduced delivery efficiency of the larger plasmid and the necessity of performing two recombination events in the dual-cassette strategy may inherently be less efficient than a single integration when using a minicircle.

**Figure 6.**
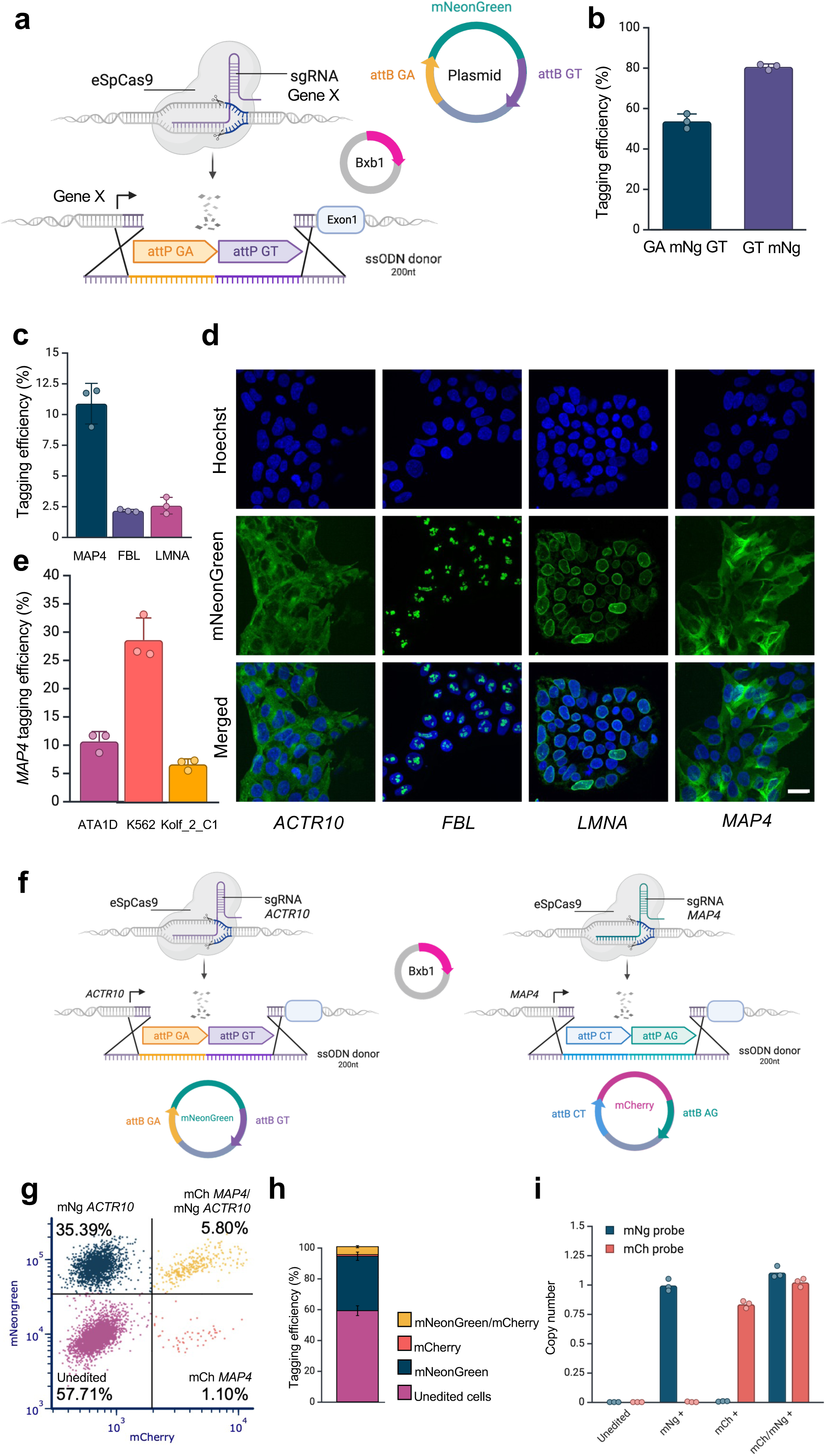
Expanding the ONE-STEP platform with dual-cassette integration and multiplexed tagging. **a.** Schematic of dual-cassette ONE-STEP tagging strategy. A single-stranded DNA HDR template (∼200 nt) containing two heterologous attP sites (attP-GA and attP-GT) and ∼50 bp homology arms was used in combination with a plasmid donor containing cargo flanked by corresponding attB variants. **b.** Bar chart comparing mNeonGreen integration efficiency at the *ACTR10* locus using the original ONE-STEP technology (∼80%) and the dual-cassette strategy (∼60%). **c.** Bar chart showing mNeonGreen integration at three additional loci (*LMNA, FBL*, and MAP4) in the A1ATD line. Tagging efficiencies ranged from 2.1% to 10.8%, depending on the locus. Data represent n=3 biological experiments; error bars indicate standard deviation. **d.** Fluorescence microscopy images confirming subcellular localization of mNeonGreen consistent with known localization patterns for all four targeted proteins (ACTR10, LMNA, FBL, MAP4). **e.** Bar chart showing extension of ONE-STEP tagging to additional cell types. Integration at the MAP4 locus resulted in tagging efficiencies of 6.6% in Kolf_2_C1 cells and 28.6% in K562 lymphoblast cells. Data represent n=3 biological experiments; error bars indicate standard deviation. **f.** Schematic of multiplexed gene integration strategy using dual-cassette donors. *ACTR10* was targeted with GA-mNeonGreen-GT, and *MAP4* with CT-mCherry-AG, in a single nucleofection. **g.** Flow cytometry analysis demonstrating successful dual tagging, with approximately 6% of cells expressing both mNeonGreen and mCherry. **h.** Bar chart quantifying dual-tagging efficiency across triplicate experiments; error bars indicate standard deviation. **i.** Double-positive and single-positive cell populations were sorted by flow cytometry, and site-specific integration at *ACTR10* and *MAP4* loci was validated by ddPCR using junction-specific probes for mNeonGreen and mCherry.

Despite the slightly reduced efficiency, the dual-cassette tagging strategy significantly enhances the versatility of our approach as we are now able to generate a universal library of donor plasmids (to be distributed via Addgene), facilitating broader adoption of this technology.

Since our technology does not require the assembly of long ssODN HDR templates but instead utilizes a short ssODN (200 nt) with approximately 50 bp of homology arms flanking the recombination cassettes, cargo integration can be easily scaled across different loci. We tested and confirmed the successful integration of mNeonGreen at three additional genomic locations (*LMNA*, *FBL*, and *MAP4*) in the hiPSC line A1ATD. Integration efficiency varied between 2.1% and 10.8%, depending on the locus (Fig. 6c). Although the efficiency was lower for some sites, by fluorescent sorting and expanding the cells we can enrich a pure population with the integrated construct. We also note that expression of the tagged genes remains stable for at least 1 month in culture.

To assess the accuracy of the tagging, we used fluorescence microscopy to compare the subcellular localization of mNeonGreen with the known localization of the tagged protein. For all four targeted loci, mNeonGreen localized as expected, confirming successful tagging (Fig. 6d).

We also extended the ONE-STEP tagging technology to additional cell types beyond the hiPSC lines (A1ATD and Kolf_2_C1), testing it in the K562 lymphoblast cell line. Integration of a 0.8 kb fluorescent tag at the *MAP4* locus yielded varying efficiencies, ranging from 6.6% in Kolf_2_C1 cells to nearly 28.6% in K562 cells (Fig. 6e).

We also leveraged the use of distinct attP sites (e.g., CT and AG) to generate a dual-cassette donor construct (CT-mCherry-AG) for a multiplexed gene integration experiment. Specifically, we aimed to simultaneously tag two different genes within the same cell: *ACTR10* with mNeonGreen using the GA-mNeonGreen-GT plasmid, and *MAP4* with mCherry using the CT-mCherry-AG plasmid (Fig. 6f).

Flow cytometry analysis confirmed successful dual tagging, with approximately 6% of cells expressing both fluorescent tags (Fig. 6g, 6h). Double-positive and single-positive cell populations were sorted, and integration at the correct genomic loci was validated by ddPCR utilising the primers that span the fluorescent tags and the endogenous loci (Fig. 6i). Double-positive cells were also imaged, revealing fluorescence signals consistent with the expected subcellular localization of ACTR10 and MAP4, further confirming correct tagging and minimal cross-talk between the GA-GT and CT-AG sites (Fig. S5).

In conclusion, the dual-cassette ONE-STEP tagging strategy represents a substantial advance in the accessibility and scalability of our genome engineering platform across different genomic loci and cell types. By removing the need for custom minicircle donor production and instead employing a short, universal ssODN in combination with pre-made plasmid donors (available from Addgene), this method enables rapid and cost-effective adaptation across a wide range of target loci and cell types. Although the tagging efficiency is slightly reduced compared to the original minicircle-based system, this trade-off is justified by the operational flexibility and convenience of this system for certain applications.

Importantly, the generation of a standardized library of dual-cassette donor plasmids, designed with orthogonal attB sites, enables multiplexed and combinatorial tagging experiments, as demonstrated by the successful dual integration of fluorescent tags at ACTR10 and MAP4. This modularity opens the door to more complex synthetic biology applications, including multiplexed lineage tracing, protein interaction mapping, and organelle-specific reporter systems, all from a single universal workflow.

## Discussion

In this study, we develop the ONE-STEP tagging platform, a streamlined, high-efficiency genome engineering system that combines CRISPR-Cas9 targeting with Bxb1 integrase-mediated site-specific DNA integration. By coupling attP site insertion via HDR with subsequent Bxb1-mediated recombination of a circular donor plasmid containing an attB site, we achieve precise transgene integration of cargo up to 14.4 kb in a single delivery step. Tagging efficiency is further enhanced through optimization of the Bxb1 integrase—using variants with improved nuclear localization and catalytic activity—and by modulating DNA repair through DNA-PK inhibition.

A recent study from Jacob Corn’s lab (50) highlighted a cautionary aspect of using DNA-PKs inhibitors such as AZD-7648. While these inhibitors are known to enhance HDR efficiency, Corn’s team reported that their use in combination with sgRNAs can lead to kilobase-to megabase-scale deletions and chromosomal translocations across multiple human cell types. These large-scale genomic alterations, typically undetectable by short-read sequencing, represent a significant fraction of editing outcomes and raise concerns for therapeutic applications, although the frequency of such events is less when the editing site is further from the telomeres and can be further reduced by co-treatment with a polQ inhibitor (polQi2). For research applications they are less problematic as they can be identified and removed during clone genotyping. Interestingly, these overlooked consequences are not limited to DSB-inducing genome editors; recent reports have associated base editing in HSPCs with translocations as well (51, 52).

In our own implementation of ONE-STEP tagging, we used shorter inhibitor exposure (24 hours vs. 72 hours) and lower AZD-7648 concentrations than those reported in the Corn study. While we have not yet assessed whether our optimized conditions produce similar genomic rearrangements, this will be important to address for therapeutic application of our method. Importantly, even without AZD-7648 treatment, ONE-STEP tagging achieves robust performance, including ∼30% efficiency at the *ACTR10* locus in hiPSCs—highlighting the system’s effectiveness without pharmacological enhancement and its potential for safer genome engineering in sensitive contexts.

To increase the practicality of the method, we developed a dual-cassette tagging strategy that eliminates the need to generate a new minicircle for each donor construct. This enables fully off-the-shelf integration using standard reagents and a modular set of cargo plasmids available via Addgene. We demonstrate the versatility of ONE-STEP tagging across a wide range of human cell types—including cancer lines, primary T cells, and notably human induced pluripotent stem cells (hiPSCs)—where we successfully integrated cargos up to ∼14 kb in size. Strikingly, at the *ACTR10* locus in hiPSCs, we achieved tagging efficiencies of up to 80%, a level of performance not reported with other technologies. ONE-STEP tagging is currently the only available method that delivers such robust, reproducible integration efficiency in hiPSCs, where alternative systems often fall short. For instance, prime editing systems such as eePASSIGE, though elegant in design, show very low tagging efficiencies in hiPSCs (typically below 1%), likely due to challenges in editor delivery and limitations in large DNA integration. Meanwhile, classical HDR approaches require labour-intensive donor plasmid preparation with long homology arms and often result in low and inconsistent efficiencies. By contrast, ONE-STEP enables rapid, high-efficiency tagging in hiPSCs using off-the-shelf, modular donor vectors, overcoming these limitations.

A key strength of the system is its compatibility with simultaneous editing at other sites; we show that gene tagging can be paired with knockout or HDR at a second locus, enabling complex, multiplex genetic engineering. We further demonstrate its translational potential in primary human T cells, where we precisely integrated a 4.4 kb EGFP-expressing construct (similar in length to a CAR) at the *TRAC* locus as well as simultaneous knockout of B2M. This would enable generation of universal, off-the-shelf CAR-T cells capable of evading alloresponsive T cell attacks in a single step using the CAR integration to select for double-edited cells. This strategy can be extended to additional edits, such as PD-1 knockout, to reduce exhaustion and improve persistence, as supported by previous findings (Eyquem et al., 2017). Finally, in contrast to prime editing approaches that are limited by the number of recombination sites that can be introduced per event, our HDR-recombinase platform readily accepts pre-assembled, unmodified dual-cassette donors, streamlining the workflow and enabling true off-the-shelf engineering.

In summary, the ONE-STEP tagging system represents a major advance in genome engineering, providing a fast, versatile, and scalable approach to precise gene integration with broad applications ranging from basic research to therapeutic cell engineering. In the future our technology could be used with orthogonal recombinase systems such as Pa01 to further enhance the adaptability and multiplexing capabilities.

## Data availability

Raw data for flow cytometry, sequencing and vector maps is available on Zenodo (5) under a CC-BY-4.0 license. Plasmids are available from Addgene.

## Acknowledgements

We would like to thank the flow cytometry facility in Cellular Operations for analysis support, Sequencing Operations for high throughput sequencing and to Ben Davies for supplying the Bxb1 and attachment site sequences. We are grateful to members of the Bassett lab for helpful comments on the manuscript, advice, discussions and support.

