## Supplementary figures S1-S5 for "ONE-STEP tagging: a versatile method for rapid site-specific integration by simultaneous reagent delivery"

Consisting of
Supplementary Figures S1-S5

### Fig S1

**a**

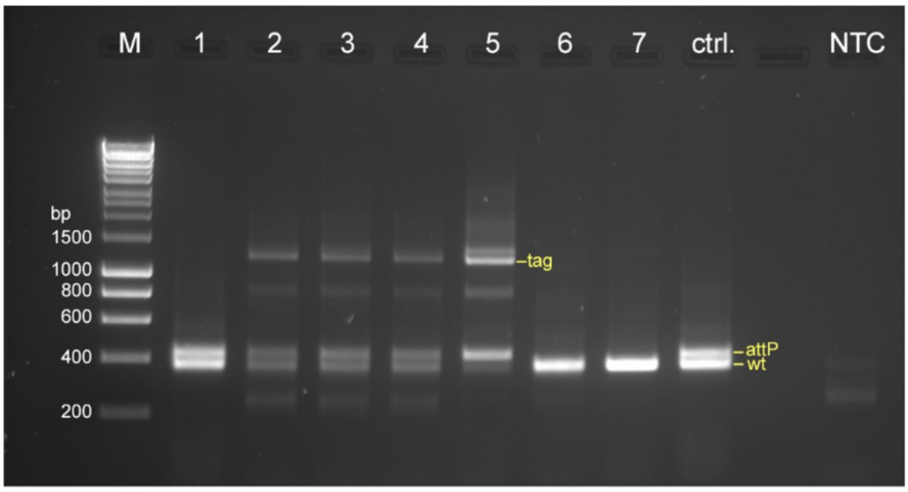


| | | | | | | | | |
|---|---|---|---|---|---|---|---|---|
| **ssODN-attP** | + | + | + | + | + | - | - | + |
| **Bxb1 plasmid** | - | 250 ng | 250 ng | 500 ng | 500 ng | 500 ng | - | - |
| **cargo-attB mNeonGreen** | - | 96 ng | 192 ng | 192 ng | 384 ng | 384 ng | - | - |

**b**

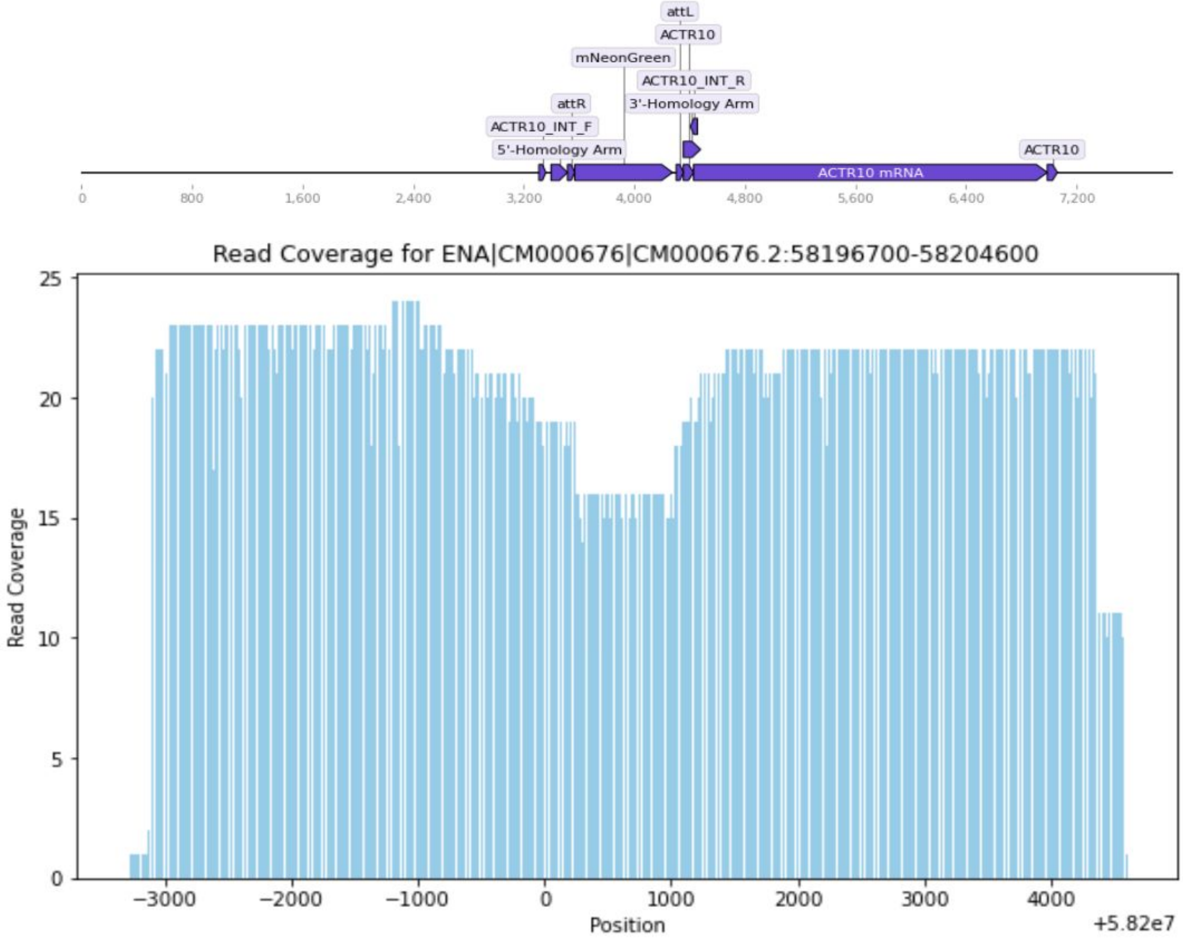


**Fig S1. PCR genotyping of mNeonGreen *ACTR10* ONE-STEP tagging**. **a.** PCR and gel electrophoresis readout of ssODN (attP) and cargo attB mNeonGreen integration (tag) at the *ACTR10* locus. The integration is analysed under different experimental conditions as well as controls lacking the ssODN, Bxb1 plasmid and cargo as indicated to validate integration events and specificity of PCR amplification. The gel image shows bands corresponding to the PCR fragments, with expected sizes indicated in yellow. Primers are annotated in Supplementary Plasmid map. The lanes represent various treatment conditions, allowing comparison of integration efficiency and specificity.
**b.** Plot of read coverage of attB mNeonGreen integration at the *ACTR10* locus - data is from Cas9-enrichment combined with Oxford Nanopore long read sequencing. The map of the integrated locus is shown above the plot. The mNeonGreen fragment is not present in all alleles, and thus there is a lower coverage over this region. The coordinates refer to the edited allele.

### Fig S2

a

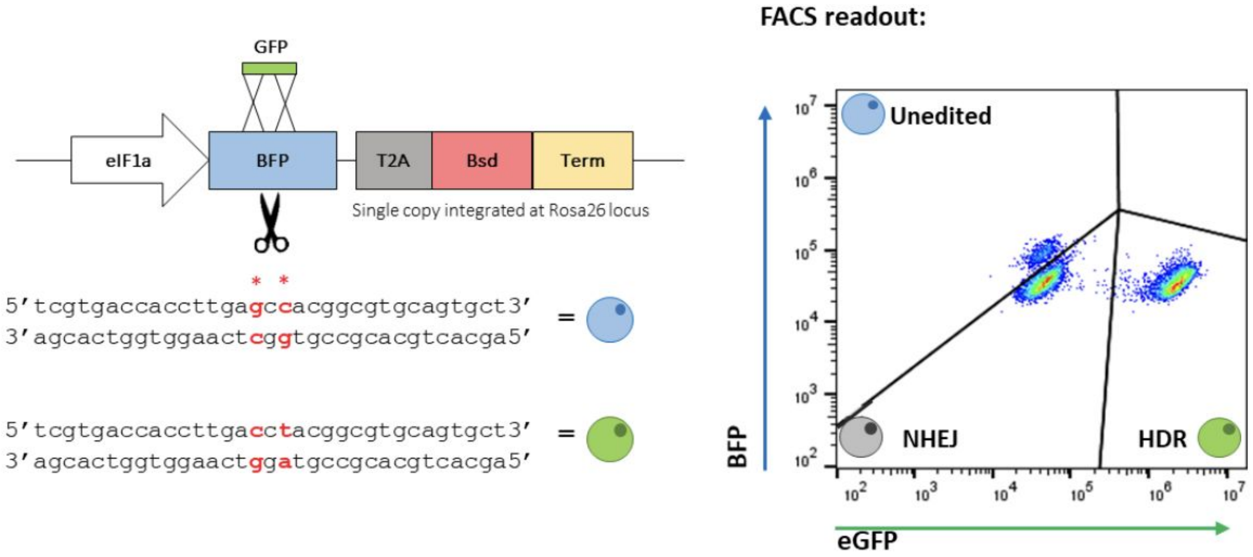


b

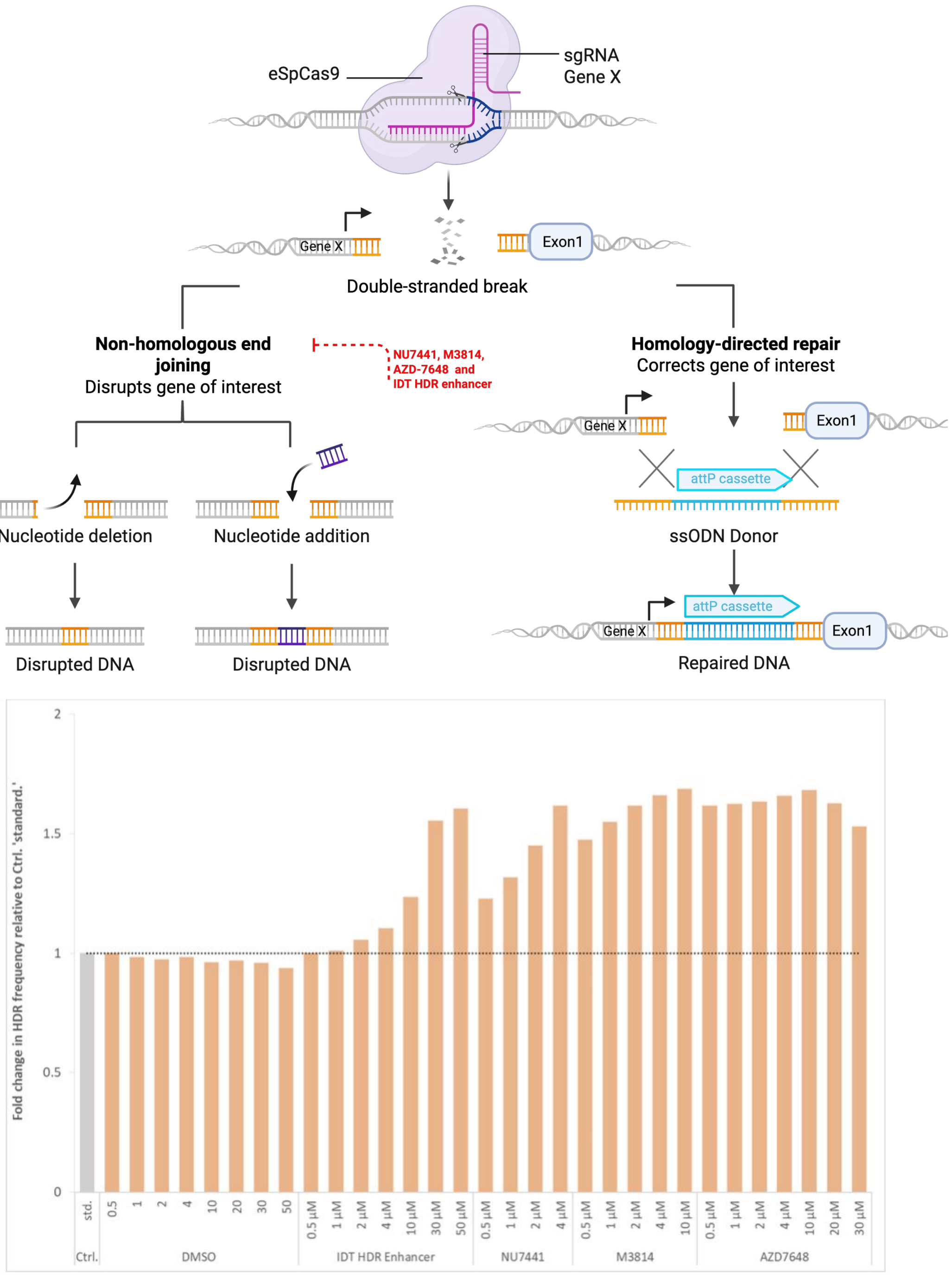


**Fig S2. Testing optimal concentration of DNAPK inhibitors. a.** Schematic of the BFP reporter system and FACS readout upon KO and HDR events. **b.** Screening to test the optimal concentration of several DNA-PK inhibitors (IDT HDR Enhancer v2, AZD-7648, NU7441, M3814) and a DMSO control followed by fold change analysis of HDR efficiency (BFP to GFP conversion) compared to control. Schematic of experiment on the top. The graph on the bottom depicts the fold change in HDR efficiency for each treatment condition, illustrating the relative effectiveness of each DNA-PK inhibitor in promoting HDR.

### Fig S3

**a**

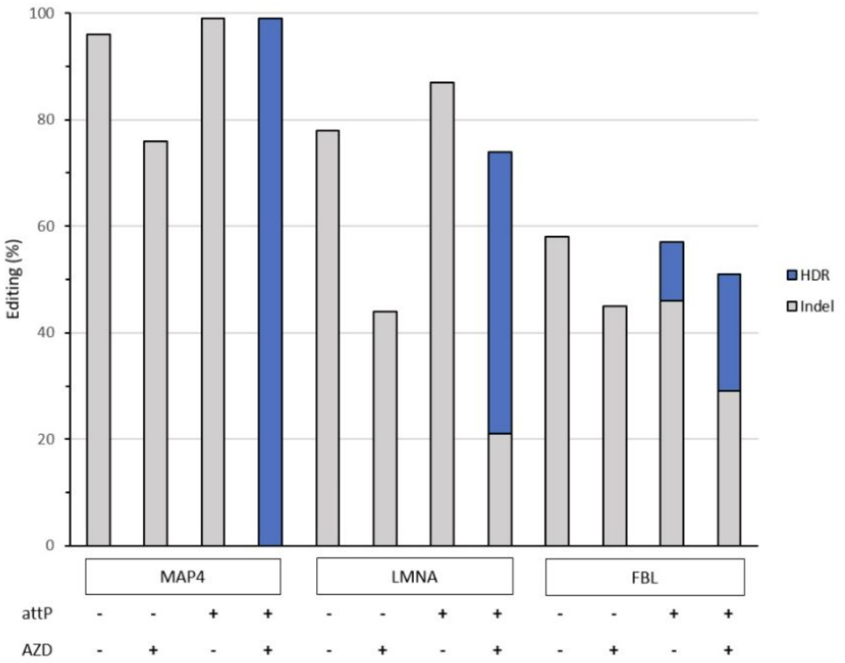

Editing (%)
100
80
60
40
20
0
HDR
Indel
MAP4
LMNA
FBL
attP
AZD


**b**

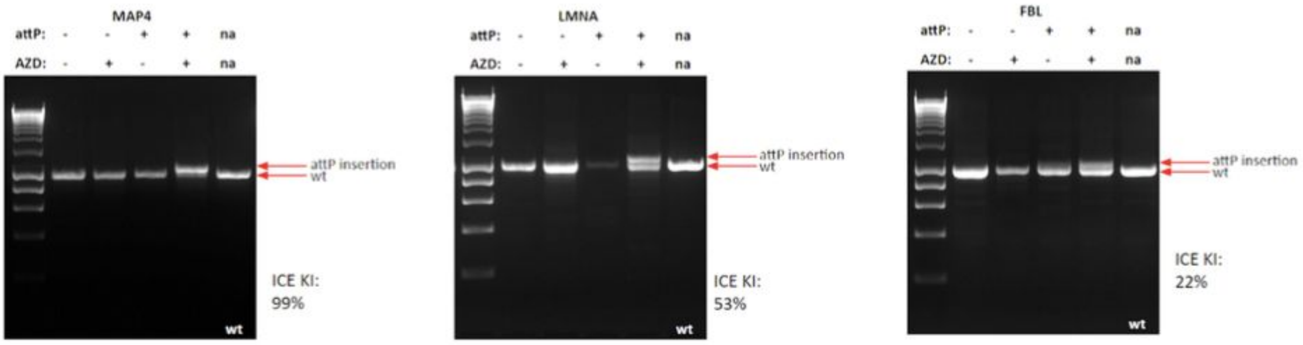

MAP4
attP:
AZD:
na
attP insertion
wt
ICE KI:
99%
LMNA
ICE KI:
53%
FBL
ICE KI:
22%


**c**

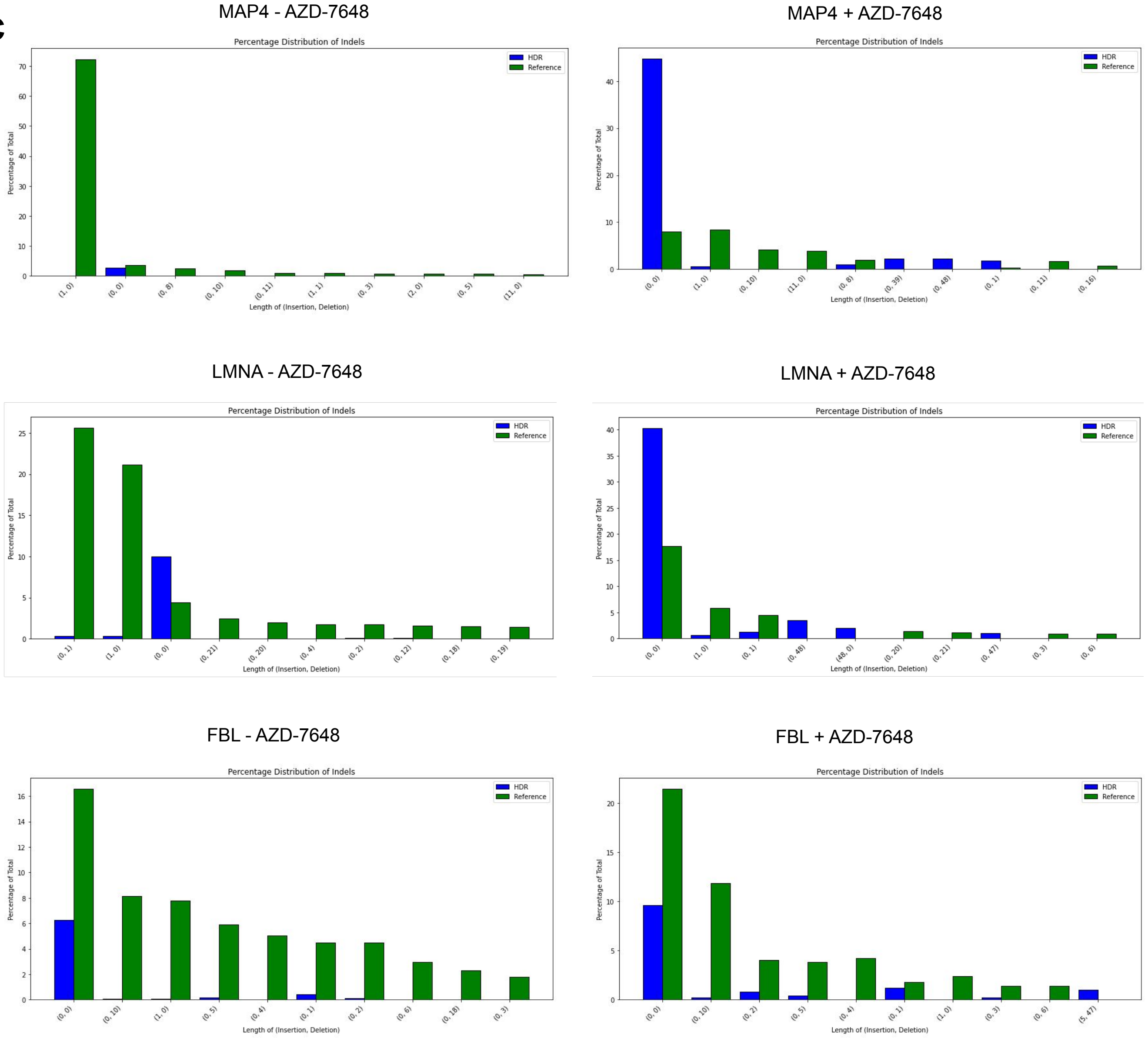

MAP4 - AZD-7648
MAP4 + AZD-7648
LMNA - AZD-7648
LMNA + AZD-7648
FBL - AZD-7648
FBL + AZD-7648
Percentage Distribution of Indels
HDR
Reference
Percentage of Total
Length of (Insertion, Deletion)

**Fig S3. Tagging of MAP4, LMNA and FBL and subsequent analysis of attP integration by Inference of CRISPR Edits (ICE) and high throughput sequencing (HTS). a.** ICE analysis showing the HDR efficiency of the attP cassette at the 5' end of *MAP4, LMNA*, and *FBL* in the presence or absence of AZD-7648 and ssODN containing the attP site. HDR efficiencies of 99%, 53%, and 22% were achieved for *MAP4, LMNA* and *FBL* respectively, in the presence of AZD-7648 and ssODN attP in the reaction. The ICE tool has a detection threshold of approximately 5%, so any tagging or indels below this level may not be reliably detected. **b.** PCR and gel electrophoresis readout of ssODN attP integration at the MAP4, LMNA, and FBL loci. The integration is analysed under different experimental conditions as well as a control. The gel image shows bands corresponding to the PCR fragments, with expected sizes indicated in red. The lanes represent various treatment conditions, allowing comparison of integration efficiency and specificity. Controls lacking the attP oligo are included to validate the integration events and the specificity of the PCR amplification. **C.** Analysis of attP integration efficiency and fidelity through MiSeq sequencing across the *MAP4, LMNA* and *FBL* loci. Green bars depict the number of reads mapping to the wild-type allele and blue bars represent the number of reads mapping to the HDR allele as analysed by CRISPResso2. Reads have been grouped together based on the number of insertions and deletions the nature of which is indicated on the x-axis (brackets). Only the 10 most frequent insertion or deletion combinations are shown.

### Fig S4

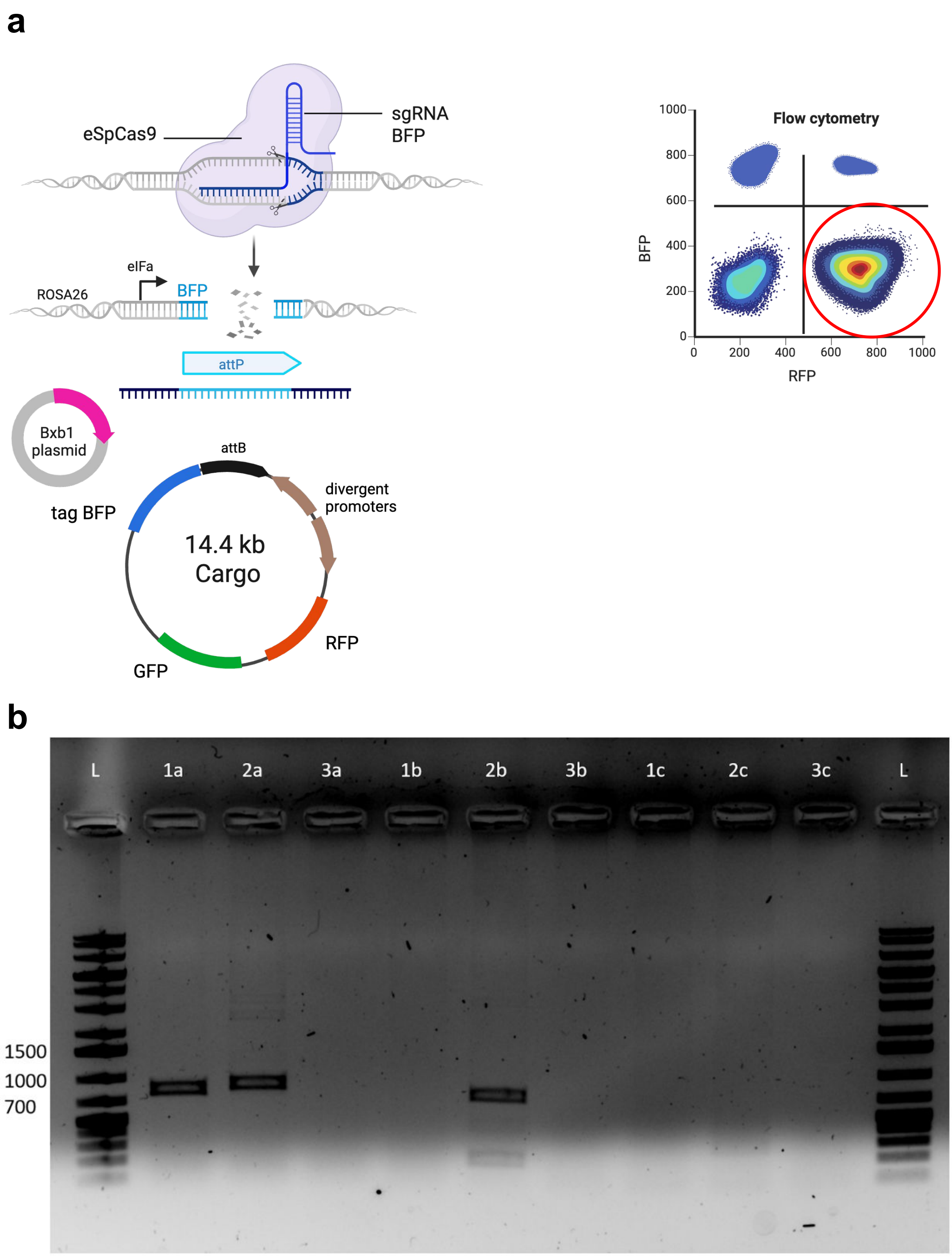


**Fig S3. ONE step tagging can integrate large cargo. a.** Schematic of reporter assay for integration experiment of a large plasmid (14.4kb) at an engineered BFP-expressing locus. A plasmid containing a set of two divergent constitutive promoters driving the expression of a tagBFP reporter gene on one side as well as RFP and GFP reporter genes on the other side is targeted for insertion into a chromosomal locus containing a constitutively expressed BFP reporter. Upon integration into the BFP-expressing genomic locus, the expression of the BFP reporter gene from the plasmid and the chromosome are both disrupted, whilst RFP and GFP expression is induced. A representative flow cytometry plot upon integration. The population highlighted in red indicates correct integration of the 14.4kb plasmid. **b.** Gel electrophoresis analysis of PCR products spanning the integration junction from insertion of the 14.4kb donor at the *BFP* locus. The PCR conditions are labelled as: a – amplification of wild-type allele (BFP_F5 and BFP_R5) – expected sizes are 796bp for the unedited allele and 854bp for the attP integration allele, b – amplification of 3' arm of integration (IG21and BFP_R5) expected size is 723bp, c – amplification of plasmid-specific product (BFP_F5 and IG21) – expected size is 270bp. The templates are labelled as 1 – 45ng of genomic DNA extraction from unedited BFP line, 2 – 45 ng of genomic DNA extraction from edited BFP line sorted for positive RFP and negative BFP, 3 – ultrapure water. L – ladder.

### Fig S5

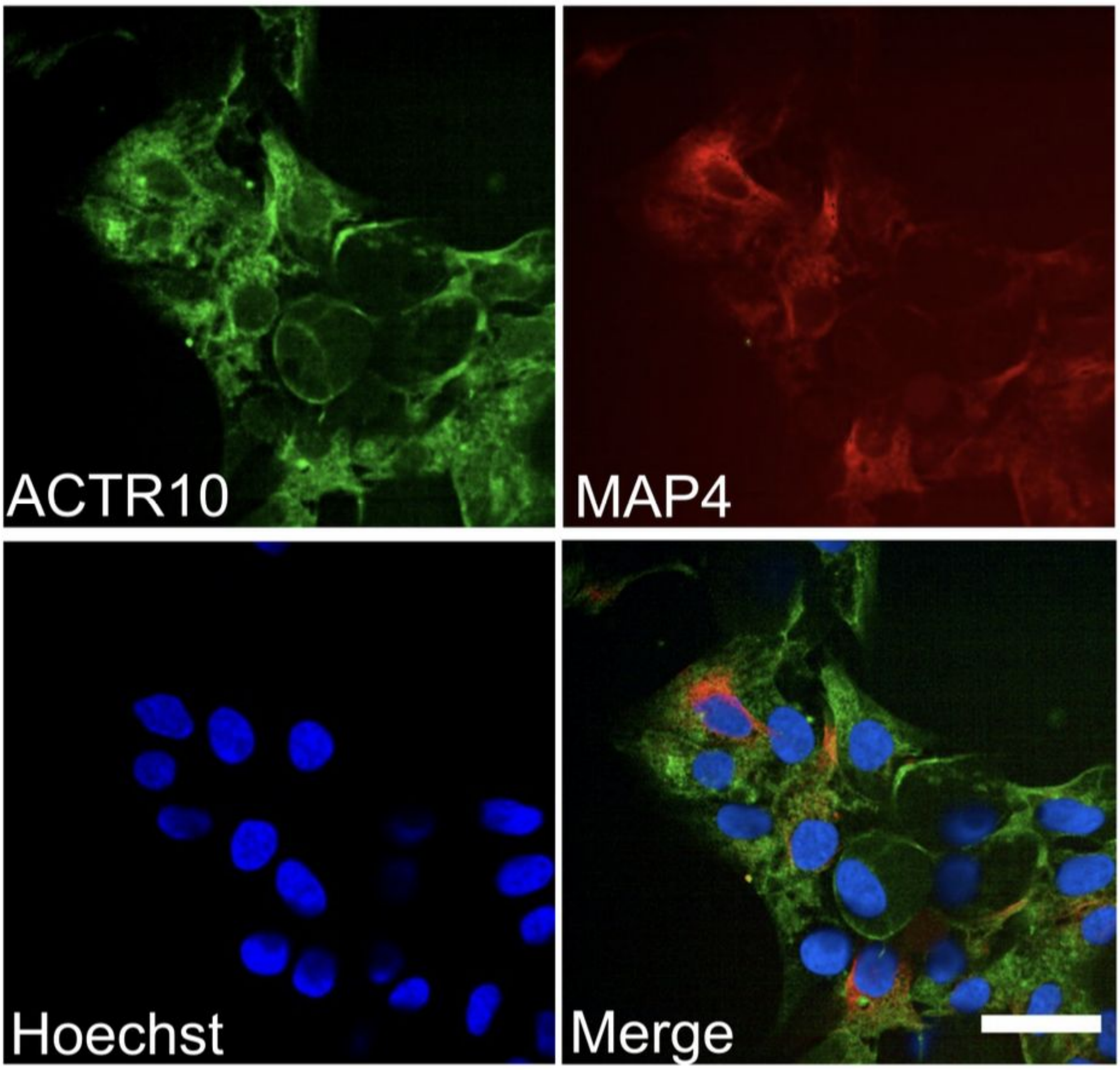


**Fig S5. Simultaneous multiplexed tagging of two different loci.** Endogenous multiplexed protein tagging with mNeonGreen and mCherry via ONE-STEP tagging at 2 loci (*ACTR10* and *MAP4*). 40x Fluorescence images of representative cells are shown. Cells have nuclear Hoechst staining (blue). Endogenous expression of mNeonGreen (green) and mCherry (red) localise according to known, location of the two tagged genes in the literature. Scale bar 30μm.
